# Sialic acid-containing glycolipids extend the receptor repertoire of Enterovirus-D68

**DOI:** 10.1101/2025.01.17.633529

**Authors:** Ashley K. Pereirinha da Silva, Jacobus P. van Trijp, Anouk Montenarie, Jelle Fok, Syriam Sooksawasdi Na Ayudhya, Roland J. Pieters, Geert-Jan Boons, Debby van Riel, Robert P. de Vries, Lisa Bauer

## Abstract

Enterovirus D68 (EV-D68) emerged as a pathogen of increasing health concern globally, particularly due to its association with outbreaks of severe respiratory diseases and acute flaccid myelitis (AFM) in children. Knowledge regarding the tissue tropism and pathogenesis of EV-D68 within the respiratory tract and central nervous system remains limited, primarily due to an incomplete understanding of the host factors that facilitate EV-D68 entry into host cells. Several cellular receptors involved in EV-D68 infections have been identified, including ICAM-5, sialylated glycoproteins, and heparan sulfate (HS). Here, we investigate the receptor requirement of a panel of EV-D68 strains covering all clades focusing on HS and sialosides utilizing glycan arrays. We found that all EV-D68 strains binding to HS harbour a cell culture adaptative substitution in the structural protein VP1 at position 271 which changes the amino acid into a positive charged one. Glycan array analyses revealed that EV-D68 strains either prefer α2,6-linked sialic acids presented on N-glycans, α2,8 linked sialic acids on gangliosides, or both. Inhibition of glycolipid biosynthesis or multivalent glycolipid mimics confirmed that ganglioside structures serve as entry receptors for certain EV-D68 strains. Lastly, we examined whether EV-D68 strains that bind to HS or glycolipids require different uncoating mechanisms. Bafilomycin A1 minimally affected cell entry of HS-binding EV-D68 strains B2/039 and B2/947 and the ganglioside preferring B1/2013 other viruses were strongly inhibited. Together, we identified that EV-D68 strains can use disialoglycolipids as novel receptors and that different EV-D68 strains show a promiscuous sialic acid binding repertoire.

## Introduction

The globally (re)emerging Enterovirus-D68 (EV-D68) is a non-enveloped, positive-sense, single stranded RNA virus (+ssRNA) and belongs to the family *Picornaviridae*, genus *Enterovirus,* species *Enterovirus deconjuncti (*formerly named *Enterovirus-D)*^1^. It is mainly associated with mild to moderate upper respiratory disease, however severe lower respiratory tract complications and neurological conditions such as encephalitis, meningitis and acute flaccid myelitis (AFM) can occur^2,3^. Until 2008, EV-D68 was considered a rare respiratory pathogen with sporadic reports worldwide^3–5^. In 2014, clusters of severe respiratory illness that coincided with an upsurge of AFM in young children^6^ were reported in the United States^7^ which then spread to Canada^8^, Europa^9^ and Asia^10–12^. Since then, EV-D68 outbreaks followed a biennial pattern between 2014-2018 that diminished to low levels during the COVID-19 pandemic^13–16^. After the COVID-19 lockdowns, infections with EV-D68 rapidly increased worldwide causing severe respiratory disease, however while other neurological complications where detected, the cases of AFM remained largely stable^13,15,17,18^. EV-D68 strains are classified into the major genotypes A through C based on VP1 sequence analysis, with genotype B subdivided into B1, B2, and B3 subclades, while genotype A splits into A1 and A2. Globally, the subclades B3 and A2 are the most prevalent EV-D68 subtypes circulating^18–20^.

Receptor usage and attachment to host cells is a critical determinant of viral entry, host range, tissue and cellular tropism as well as pathogenesis. The proteins ICAM-5^21^ and MFSD6^22^, different glycan structures such as α2,3-, and α2,6-linked sialic acids (SIA)^23–25^ and heparan sulfate (HS)^26,27^ have been identified to play a role in EV-D68 entry though their physiological relevance remains unclear^28^. ICAM-5 has shown to be relevant in airway cells, but its expression in the brain is restricted to neurons of the telencephalon^21,29^. MFSD6 was identified as receptor important for entry into respiratory cells^22^. In the brain, the expression seems to be restricted to excitatory neurons^30^. Currently it is unclear whether MFSD6 is a functional receptor contributing to the neurotropism. EV-D68 shows a preference for α2,6-linked sialic acids^24^, which are found on ciliated epithelial cells of the airways (nasal cavity, nasopharynx, oropharynx, trachea, bronchi and bronchioles). Two human sialyltransferases can synthesize α2,6-linked SIAs on terminal galactose residues: ST6GI, which is ubiquitously expressed, and ST6GII, which resides mostly in the brain^31,32^. While O-glycans mainly contain α2,3-linked sialic acids except for α2,6-linked sialic acids linked to GalNAc^33^; lipid-linked (sialo)glycan species are ubiquitously present in the central nervous system and uniquely display α2,8-linked sialic acids^34^, though their potential as enterovirus receptors remains unstudied. Heparan sulfate (HS) represents yet another class of glycosylation^35^, is omnipresent, predominantly at the basolateral side in the respiratory tract^36^ and ubiquitously displayed in the CNS^37^, though its binding is currently linked mainly to cell culture adaptations^26,27^. Despite the promiscuous receptor usage of EV-D68, it is unclear which receptors contribute to the respiratory and neurotropism. Viral attachment to a host receptor on the cell surface initiates receptor-mediated endocytosis, where uncoating is triggered when an uncoating receptor binds to the GH loop of VP1, displacing the pocket factor from a hydrophobic pocket within the viral capsid^38^. While most enteroviruses only require receptor-mediated uncoating, EV-A71 and EV-D68 additionally require endosomal acidification, with rhinovirus A2 being unique in relying solely on acidification without receptor interaction for uncoating^39^. The functional importance and the molecular consequences of HS and SIA binding to the virus particle remain to be fully elucidated.

In this study, we set out to investigate the receptor specificity of EV-D68 strains from different clades using glycan arrays containing either HS or N-glycans and glycolipid structures terminating in α2,3/α2,6/α2,8-linked sialosides. Furthermore, we investigate whether specific receptor requirements of EV-D68 strains is linked to differences in their uncoating strategies.

## Results

### Enterovirus-D68 strains with a positively charged amino acid in the structural protein VP1 position 271 use heparan sulfate as entry receptors

Previously, it has been shown that the EVD68 strains B2/947 and B3/1013 bind to heparan sulfate (HS)^26,27^. In both cases the amino acid substitution E271K in VP1 was identified as responsible for HS recognition. Here, we analysed whether EV-D68 strains derived from all clades according to a phylogenetic tree derived from platform Nextstrain^40^ (Figure 1A) are able to bind to HS using a previously published glycan array^41^ containing almost a 100 unique di-, tetra-, hexa-, and octasaccharides differing in backbone composition and sulfation pattern. We included the known HS binders EV-D68 strains B2/947 and B3/1013 as positive controls and the Fermon strain as negative control as this virus does not bind to HS^26^. To evaluate the binding interactions of infectious EV-D68 strains with various HS present on the array, viral detection was performed by using a primary antibody targeting the structural protein VP1, followed by visualization with an Alexa-647-conjugated secondary antibody. Strong fluorescence intensity indicative of HS binding was observed for the EV-D68 strains B2/039 and B3/1013 but only minor intensity was observed for the strains B2/947, A/2012, A2/2018, B1/2013, B3/2019 and Fermon (Figure 1B). Sanger sequencing of all used EV-D68 virus stocks revealed two amino acid substitutions in the EV-strain B2/039 (VP1 E271R, VP3 T236K), two substitutions in A2/2018 (VP1 R255K and K270Y) and a single substitution in the EV-D68 strain A/2012 (VP1 T256N) (Table 1) compared to the sequences of the original strain. Similar to the E271K VP1 cell culture adaptation found in B2/947 and B3/1013^27^, EV-D68 strain B2/039 contains a charge reversion in this position (E271R), while all other strains have retained a negatively/neutrally charged residue at this position.

**Figure 1.**
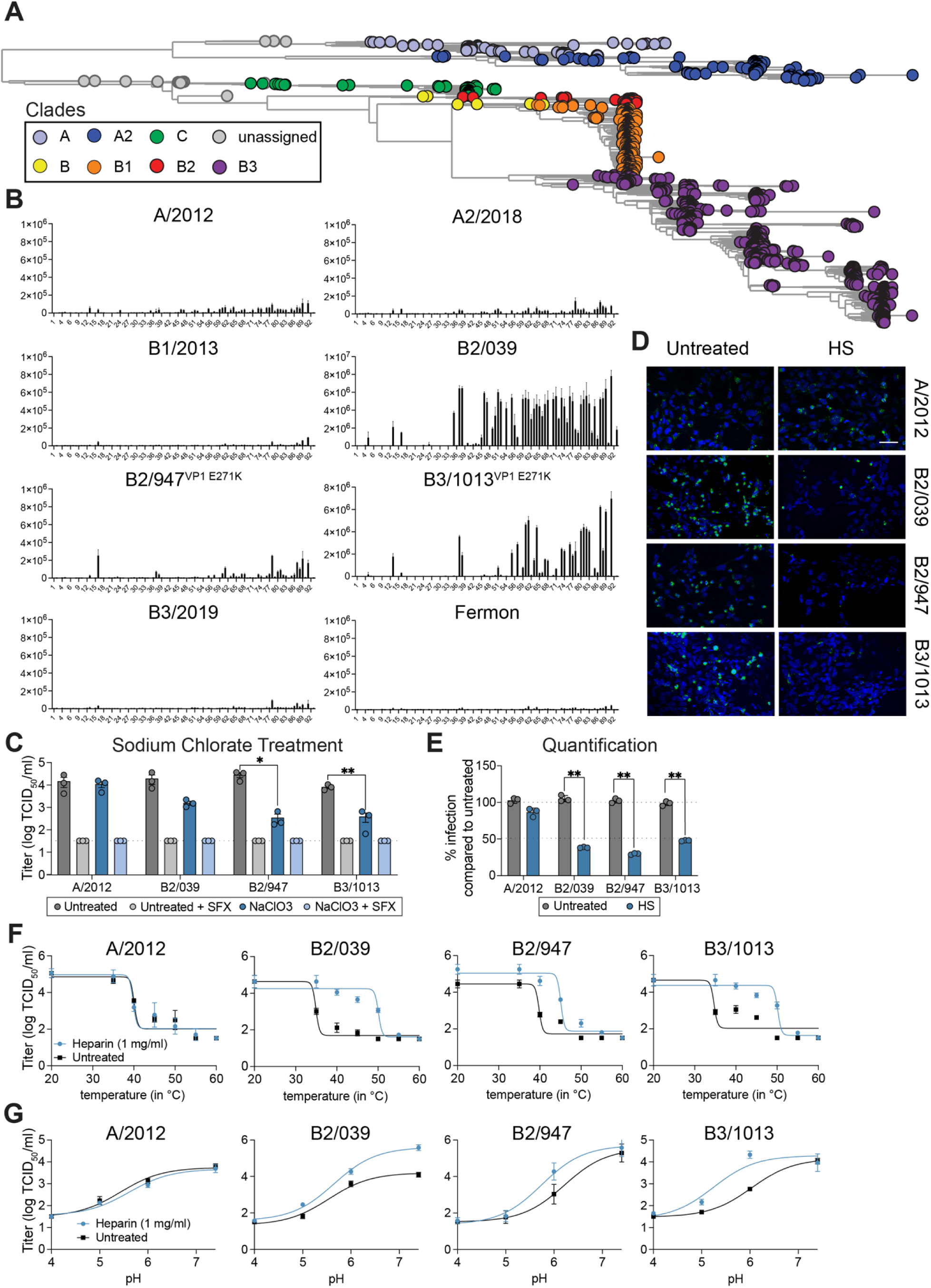
Enterovirus-D68 strains carrying a VP1 mutation bind to sulfated glycosaminoglycans. (A) Phylogenetic tree of the EV-D68 clades derived from the Nextstrain database^40^ (B) Heparin sulfate binding profile of various Enterovirus-D68 (EV-D68) strains. Two independent arrays from two different batches of virus stocks were performed. Data displayed represent one independent experiment. (C) In a single-cycle viral replication assay, RD cells were pre-treated with NaClO_3_ and infected with EV-D68 strains at multiplicity of infection (MOI) of 1. 10µM of the replication inhibitor (*S*)-fluoxetine (SFX) targeting the viral protein 2C was used as positive control. 24 hours post infection, virus titers of cell lysates were determined by endpoint dilution. Data represent values from averaged technical replicates from three independent experiments ± the standard error of mean (SEM). Statistical analysis was performed with Student’s t-test. (D) EV-D68 strains incubated with 1mg/mL soluble heparin were used to infect RD cells at a multiplicity of infection (MOI) of 1. Cells were fixed 24 hours post infection and stained for presence of double stranded RNA (green) and nuclei were visualized with Hoechst (blue). Scale bar 50 µm. (E) Quantification of infection percentage compared to untreated EV-D68 infected RD cells. Data represent values from averaged technical replicates from three independent experiments ± SEM. Statistical analysis was performed with Student’s t-test. (F) Enterovirus-D68 strains incubated with 1mg/mL soluble heparin were exposure to ascending temperatures for 15 minutes and infectivity was assessed by endpoint dilution. Data represent mean values ± SEM from three independent experiment performed in technical triplicates. (G) EV-D68 strains incubated with 1mg/mL soluble heparin were exposure to acidic pH by carefully titrating HCl. Infectivity was assessed with endpoint dilution. Data represent mean values ± SEM from three independent experiment performed in technical triplicates. Asterisks indicate statistically significant differences P > 0.05; * P ≤ 0.05; ** P ≤ 0.01; *** P ≤ 0.001; **** P ≤ 0.0001 EV-D68, Enterovirus-D68; MOI, multiplicity of infection; SFX, (*S*)-fluoxetine; Standard error of mean, SEM;

**Table 1.**
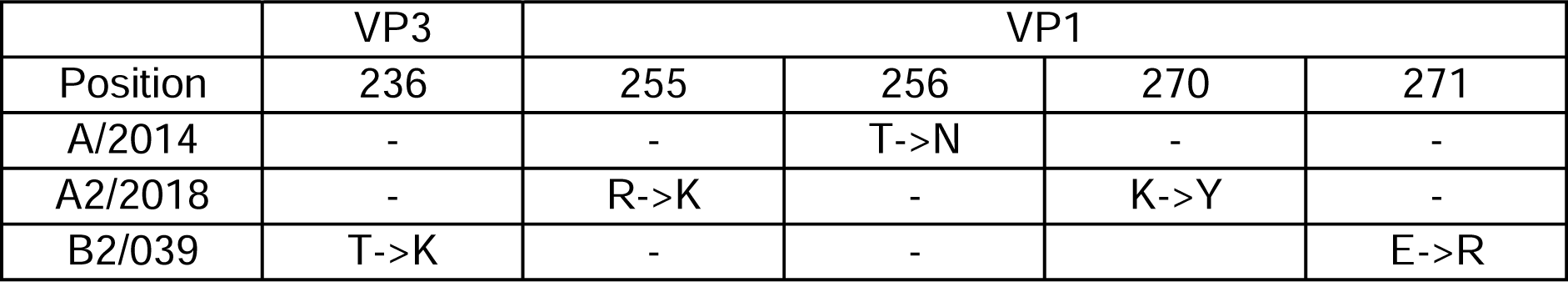
Full genome sequencing of virus stocks and identified amino acid polymorphisms.

|  | VP3 | VP1 |  |  |  |
| --- | --- | --- | --- | --- | --- |
| Position | 236 | 255 | 256 | 270 | 271 |
| A/2014 | - | - | T->N | - | - |
| A2/2018 | - | R->K | - | K->Y | - |
| B2/039 | T->K | - | - |  | E->R |

To validate the relevance of HS binding for EV-D68 replication, we pretreated different cell lines with NaClO_3_ as previously described^26,27^ to prevent cell surface sulfation^42^ and performed infection experiments. We used two strains that displayed highest fluorescent intensity towards HS namely B2/039, B3/1013 and one strain A/2012 which showed barely any fluorescent intensity on the glycan array, while the known HS-binding virus B2/947 was used as positive control. The replication inhibitor (*S*)-fluoxetine (SFX) that targets the viral protein 2C directly was used as a positive control for viral inhibition^43,44^. NaClO_3_ significantly reduced the virus titers of the EV-D68 strains B2/947 and B3/1013 but not B2/039, even though the virus titer upon NaClO_3_ treatment was one log lower at 24 hpi (Figure 1C). Similarly, NaClO_3_ treatment of SK-N-SH and A549 cells resulted in a reduction of the virus titers of EV-D68 strains B2/947 and B3/1013 (Supplement Figure 1A and 1B) excluding a cell line-dependent effect. We also observed that B2/039 showed in both cases reduced titers (one log), however only significantly reduction in A549 cells. Next, to further substantiate the interaction of B2/039, B2/947 and B3/1013 to HS, we performed neutralization experiments with soluble heparin. Virus incubation with soluble heparin significantly reduced infectivity evident by reduced intracellular double stranded RNA (dsRNA) staining, a marker for active viral genome replication (Figure 1D). Quantification of dsRNA signal showed that the infection percentage of B2/947, B3/1013 as well as B2/039 was reduced, while A/2012 appeared unaffected (Figure 1D and 1E). Lastly, to evaluate whether HS binding contributes to the stabilization of the viral capsid, we pretreated EV-D68 strains with soluble heparin and exposed the viruses to increasing temperatures and decreasing pH levels. The EV-D68 strains B2/039, B2/947, and B3/1013 but not A/2012 showed an increased thermostability (Figure 1F) and pH stability upon heparin addition (Figure 1G). Collectively, these data suggest that HS are relevant entry receptors for EV-D68 strains carrying a positively charged amino acid in the structural protein VP1 at position 271.

### Glycolipids can serve as functional Enterovirus-D68 receptors

Terminal α2,3, and α2,6-linked SIAs function as receptors for EV-D68^23–25^. To identify the sialoside receptor specificity of EV-D68 strains, we employed a glycan array containing α2,3-linked, α2,6-linked N-linked glycans and ganglioside structures containing α2,3- or α2,8-linked SIAs (Figure 2A) in a similar fashion as described previously^45^. Overall, we observed three distinct phenotypes among the EV-D68 strains: preferential binding to α2,6-linked N-linked glycans (A/2012, B2/039 and Fermon), or ganglioside structures (A2/2018) or a dual preference for α2,6 N-linked glycans and ganglioside structures (B1/2013, B2/947, B3/1013, B3/2019) (Figure 2B). Strong fluorescence intensity for ganglioside oligosaccharides was observed on multiple spots in the glycan array and correspond to the disialylated gangliosides: GT1a (E), GD3 (F), GD1c (L), GQ1b (M) which have an α2,8-linked SIA at the penultimate SIA residue. Fluorescence intensity of EV-D68 strains A/2012, B2/039, B2/947, B3/1013 and B3/2019 for α2,6-linked SIAs was highest with either 2 or 3 Lac NAc repeats, similarly as observed for human influenza A viruses^46^. Binding to α2,8-linked SIA is not a clade specific phenotype as several EV-D68 strains from multiple clades showed binding (Figure 2B). Furthermore, we have no evidence that the binding to α 2,8-linked SIA was related to cell culture adaptation since viruses without the VP1 mutation were able to bind too (Table 1). The amino acids responsible for coordination of α2,3-(Protein Data Bank [PDP]: 5BNP) or α2-6-linked (PDB: 6CVB) SIAs are located at the VP1:VP3 interface (Supplement Figure 3 and Supplement Figure 4). We aligned the capsid proteins of the sequenced EV-D68 virus stocks which showed that the amino acids coordinating SIA do not show any changes that could be linked to glycolipid binding. This indicates that EV-D68 is promiscuous for SIA binding, however whether defined amino acids contribute to glycolipid binding remains elusive.

**Figure 2.**
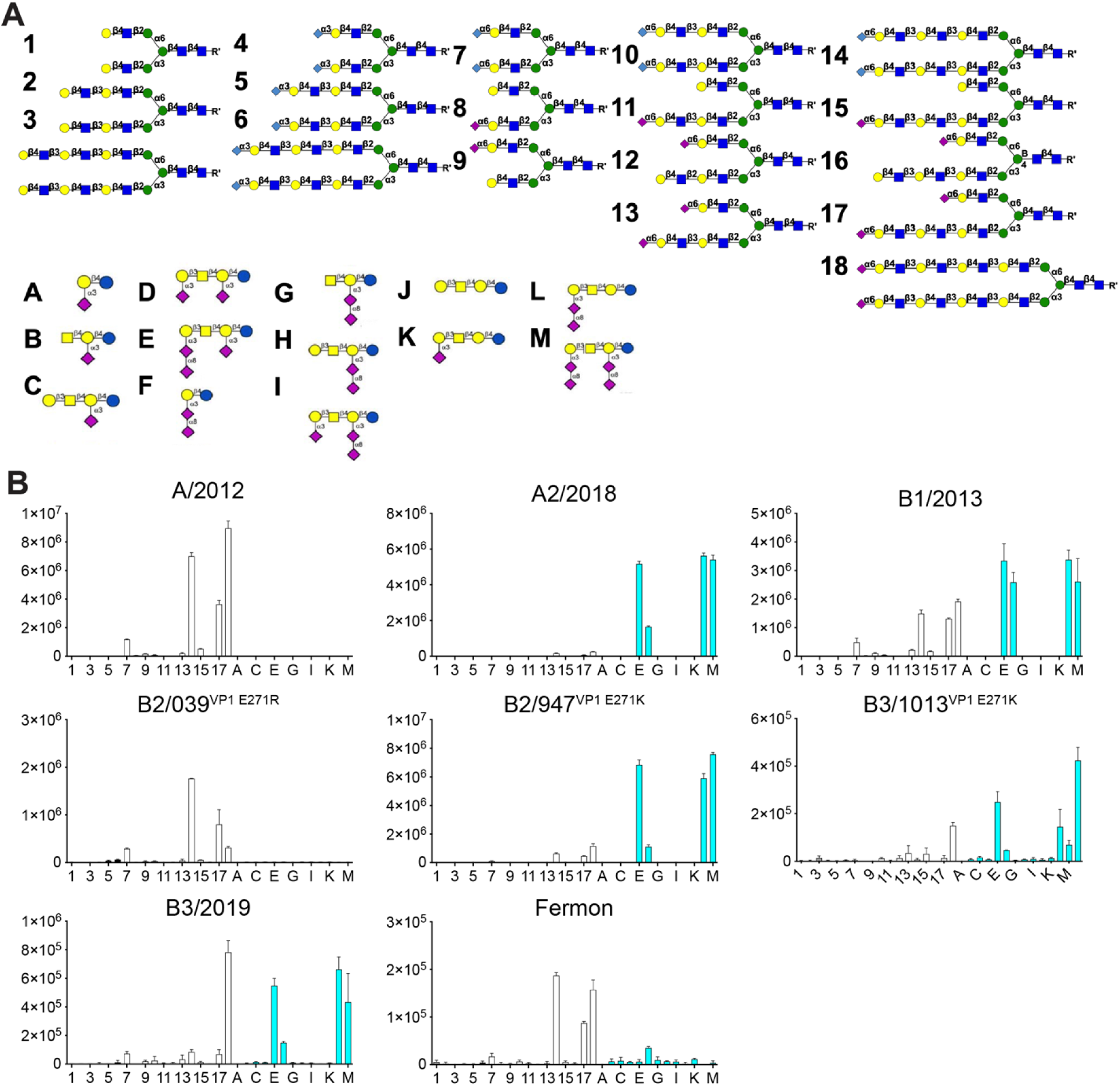
Enterovirus-D68 strains bind to SIA containing glycolipids. (A) Glycan structures containing α2,3-linked (black bars position 1-6), α2,6-linked N-linked glycans (white bars position 7-18) and ganglioside structures (cyan bars position A-N) containing α2,3- or α2,8-linked sialic acids used in the glycan array (B) N-Glycan and glycolipid binding profile of various Enterovirus-D68 strains. Two independent arrays from two different batches of virus stocks were performed. Data displayed represent one independent experiment.

To validate that glycolipids can serve as receptors, we investigated whether depletion of glycolipid biosynthesis results in a reduction in infectivity. For this we employed GENZ-123346, an inhibitor of UDP-glucose ceramide glycosyltransferase, an enzyme that catalyses the first glycosylation step in the biosynthesis of glycosphingolipids. To remove terminal α2,3-, α2,6- and α2,8-linked SIAs on glycoproteins and glycolipids we used neuraminidase (NA) derived from *Arthrobacter ureafaciens.* As a positive control, we used the virus replication inhibitor SFX. We confirmed the depletion of glycolipids through pharmacological inhibition with GENZ-123346 by measuring the reduction of the glycolipid GM1 that is specifically bound by fluorescently labelled cholera toxin by flow cytometry (Supplement Figure 2A and Supplement Figure 2B). Infection of glycolipid depleted RD cells showed a significant decrease in virus titers of EV-D68 strains that show an affinity for gangliosides (A2/2018 and B3/2019) (Figure 3A). The glycan array showed that all viruses bind to some extent to SIA, therefore NA treatment was able to reduce the virus titer of all viruses to a similar extent as the positive control SFX. These observations were validated in A459 and SK-N-SH cells (Supplement Figure 2C). Next, we performed neutralization experiments with synthesized multivalent ligands containing either GM3- or GD3 structures as inhibitors against EV-D68 strain A/2012, A2/2018, and B3/2019. EV-D68 strains were pre-incubated with GM3 and GD3 ligands or their corresponding backbone structure (BB, LAK). Subsequent infection of RD cells showed that the virus titer of EV-D68 strain B3/2019 was reduced in a dose dependent manner upon pre-incubation with the GM3 and GD3 ligands (Figure 3B), but not when incubated with their unsialylated counterparts such as the backbone structure (BB) or lactose (LAK). The inhibitory effect was less pronounced when EV-D68 strain A2/2018 was pre-incubated with the GD3 ligand. The EV-D68 strain B3/2019 also showed a significant inhibition when incubated with the backbone structure, suggesting that not only the terminal SIA seems important for binding. Together, this demonstrates novel evidence that EV-D68 strains can utilize α2,8-linked terminal SIA as entry factors. In summary, the EV-D68 strains show promiscuity in their preference towards terminal SIA structures.

**Figure 3.**
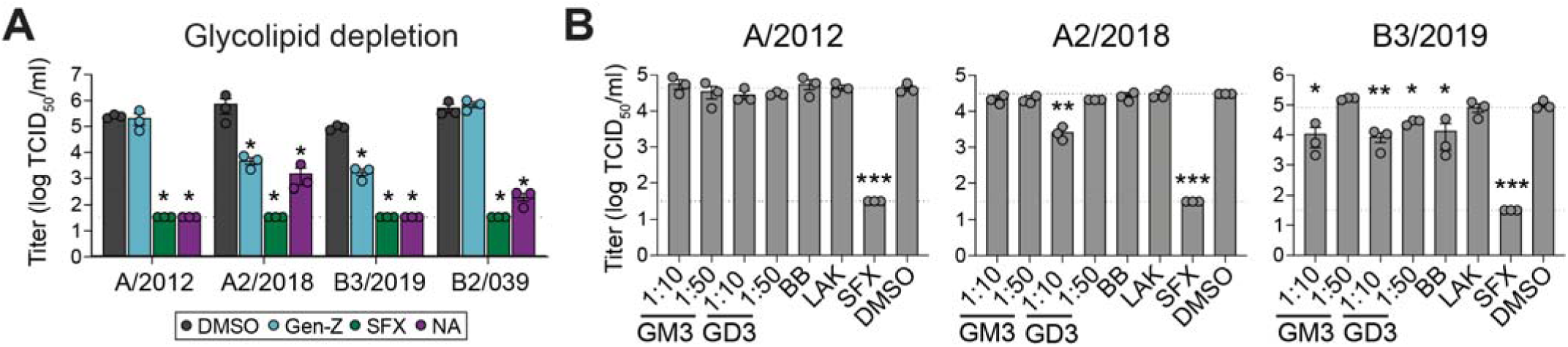
Glycolipid depletion reduces Enterovirus-D68 replication. (A) In a single-cycle viral replication assay, RD cells were first pre-treated with 5µM GEN-Z123346 (Gen-Z) for 72hrs or 100 mU/mL *Arthrobacter ureafaciens* neuraminidase (NA) for 2hours. After pretreatment, RD cells were infected with Enterovirus-D68 (EV-D68) strains at multiplicity of infection (MOI) of 1. 10µM of the replication inhibitor (*S*)-fluoxetine (SFX) targeting the viral protein 2C was used as positive control. 24 hours post infection, virus titers of lysates were determined by endpoint dilution. Data represent values from all technical replicates ± the standard error of mean (SEM). Data represent values from averaged technical replicates from three independent experiments ± the standard error of mean (SEM). Statistical analysis was performed with One Way Anova comparing the values to mock treatment. (B) EV-D68 strains were pre-incubated with the multivalent glycolipid mimics GM3 and GD3 and their corresponding backbone (BB, LAK) for 1 hours at 33°C. Afterwards RD cells were infected with MOI 1 and 24 hours post infection virus titers of cell lysates were determined by endpoint dilution. Data represent values from averaged technical replicates from three independent experiments ± SEM. Statistical analysis was performed with One Way Anova comparing the values to mock treatment. Asterisks indicate statistically significant differences P > 0.05; * P ≤ 0.05; ** P ≤ 0.01; *** P ≤ 0.001; **** P ≤ 0.0001 GenZ, GEN-Z123346; NA, neuraminidase; EV-D68, Enterovirus-D68; MOI, multiplicity of infection; SFX, (*S*)-fluoxetine; Standard error of mean, SEM; BB, backbone structure; LAK, laktose

### Enterovirus-D68 strains with varying receptor specificities show similar capsid stability profiles

It has been shown that EV-A71 HS binding comes with a trade-off for capsid stability^47^. After identifying that EV-D68 strains have an affinity for HS and/or glycolipids, we wanted to understand whether strains with different receptor repertoires also differ in their capsid stability. Therefore, we assessed the infectivity of different EV-D68 strains under changing temperature and pH conditions. We compared the EV-D68 strains to the well-studied lab-adapted Fermon strain. All HS-binding strains demonstrated significantly lower pH stability compared to the Fermon strain (Figure 4A). Among the other EV-D68 strains, only B1/2013 was significantly less pH stable compared to the control Fermon strain (Figure 4B). Thermostability experiments showed that the HS-binding strains B2/947 and B2/039, but not B3/1013 were significantly less stable compared to the Fermon strain (Figure 4C). Among the other EV-D68 strains, B1/2013 and B3/2019 exhibited a significant reduction in thermostability compared to Fermon (Figure 4D). Although not significant, the EV-D68 strain A/2012 showed a trend of higher temperature stability compared to Fermon. Together, these data show that there are no large differences in the pH sensitivity or thermostability of viruses preferentially binding to HS or other types of SIA receptors.

**Figure 4.**
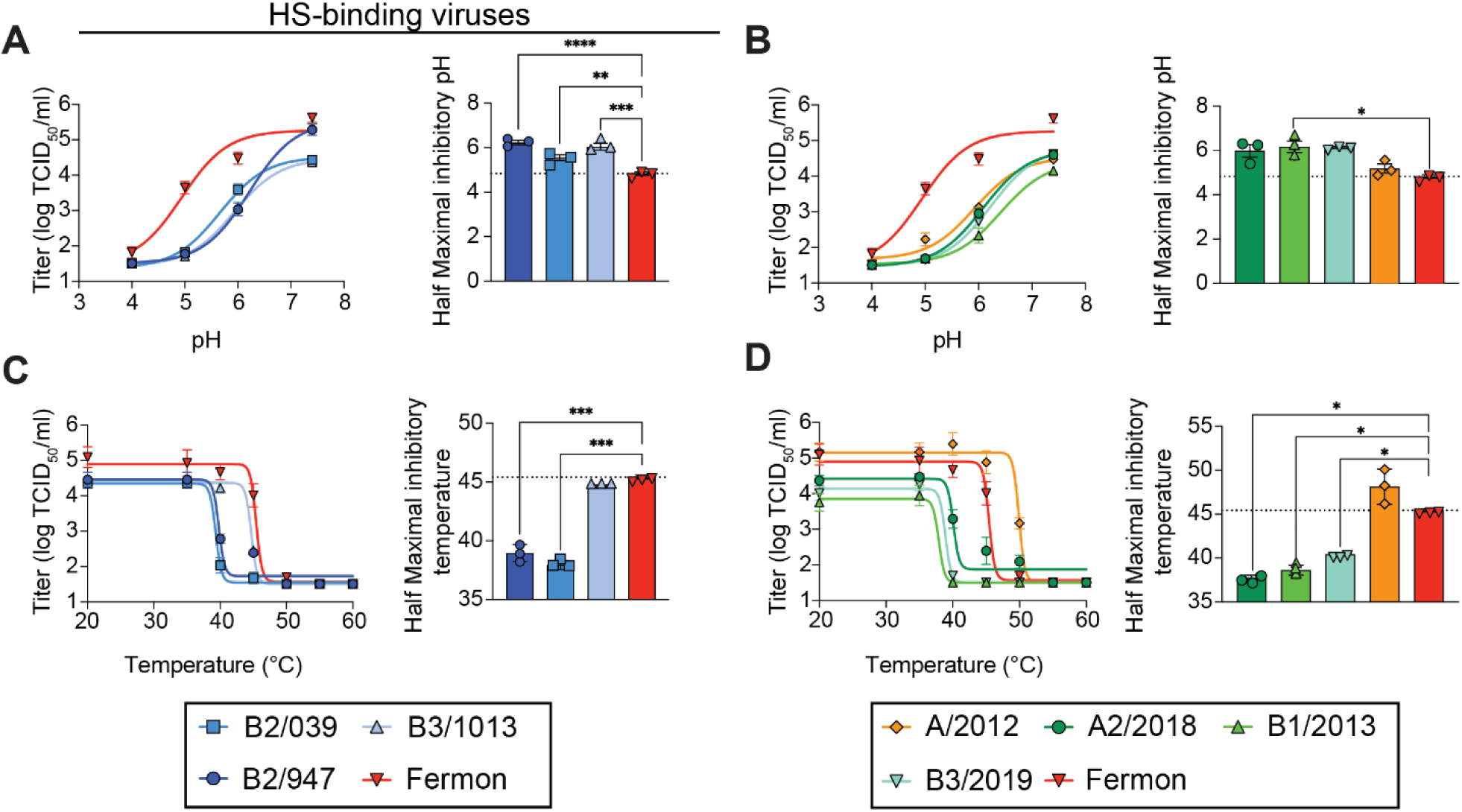
Variation in pH and thermostability stability among Enterovirus-D68 strains. (A) Heparan Sulfate (HS)-binding or (B) other Enterovirus-D68 (EV-D68) strains were incubated at different pH and infectivity was assessed by endpoint dilution. Sigmoid Curve fit was used to calculate the 50% inhibitory pH value (IC50). Data displayed represents averages of technical replicates ± standard error of the mean (SEM) and are derived from three independent experiments. Acid stability of EV-D68 strains were compared to the lab adapted Fermon strain and statistical significance was analysed by One-way ANOVA with the Tukey post hoc test test. (C) HS-binding EV-D68 strains or (D) other EV-D68 strains were incubated at different temperatures ranging from 20°C to 60°C for 15 minutes. Infectivity was assessed by endpoint dilution. Sigmoid Curve fit was used to calculate the IC50 temperature values. Data displayed represents averages of technical replicates ± SEM and are derived from three independent experiments. Thermostability of EV-D68 strains compared to the lab adapted Fermon strain was analysed by One-way ANOVA with the Tukey post hoc test test. Asterisks indicate statistically significant differences P > 0.05; * P ≤ 0.05; ** P ≤ 0.01; *** P ≤ 0.001; **** P ≤ 0.0001. HS, sulfated glycosaminoglycan; EV-D68 Enterovirus-D68, 50% inhibitory concentration, IC50; SEM, standard error of the mean

### Enterovirus-D68 strains vary in their acid dependency for entry independent of their receptor preference

Enteroviruses can use receptor mediated uncoating or like EV-D68 require additionally acidification within the endosome for their genome release^28^. Additionally, it has been shown that the lab-adapted HS-binding EV-D68 strain B2/947 uncoats in an acid-independent way^26^. Thus, we wanted to explore whether acid-independent uncoating is a hallmark of HS-binding EV-D68 strains and whether other EV-D68 viruses also require different uncoating cues. Therefore, we compared the sensitivity of EV-D68 strains towards the lysosomotropic V-ATPase inhibitor bafilomycin A1 (BAF-A1). Infection of cells pre-treated with BAF-A1 showed that the virus production of HS-binding strains B2/039 and B2/947 was hardly inhibited, whereas B3/1013 was (Figure 5A). The virus replication of all other strains EV-D68 except B1/2013 was reduced upon BAF-A1 treatment. Quantification of the infection percentage of untreated or BAF-A1 treated cells revealed similar virus production levels. The HS-binding strains B2/039 and B2/947 but not B3/1013 showed similar infection percentages whereas all other EV-D68 strains except B1/2013 showed a reduction in infection percentage upon BAF-A1 treatment (Figure 5B and Figure 5C). This likely indicates that that not all HS-binding EV-D68 strains uncoat in an acid-independent way. Other EV-D68 strains rely on endosomal acidification during uncoating similar to the positive control EV-D68 strain Fermon and RV-B14. Furthermore, it suggests that the HS-binding strains B2/039 and B2/947 hardly require endosomal acidification but rather uncoat through destabilization initiated by HS receptor engagement.

**Figure 5.**
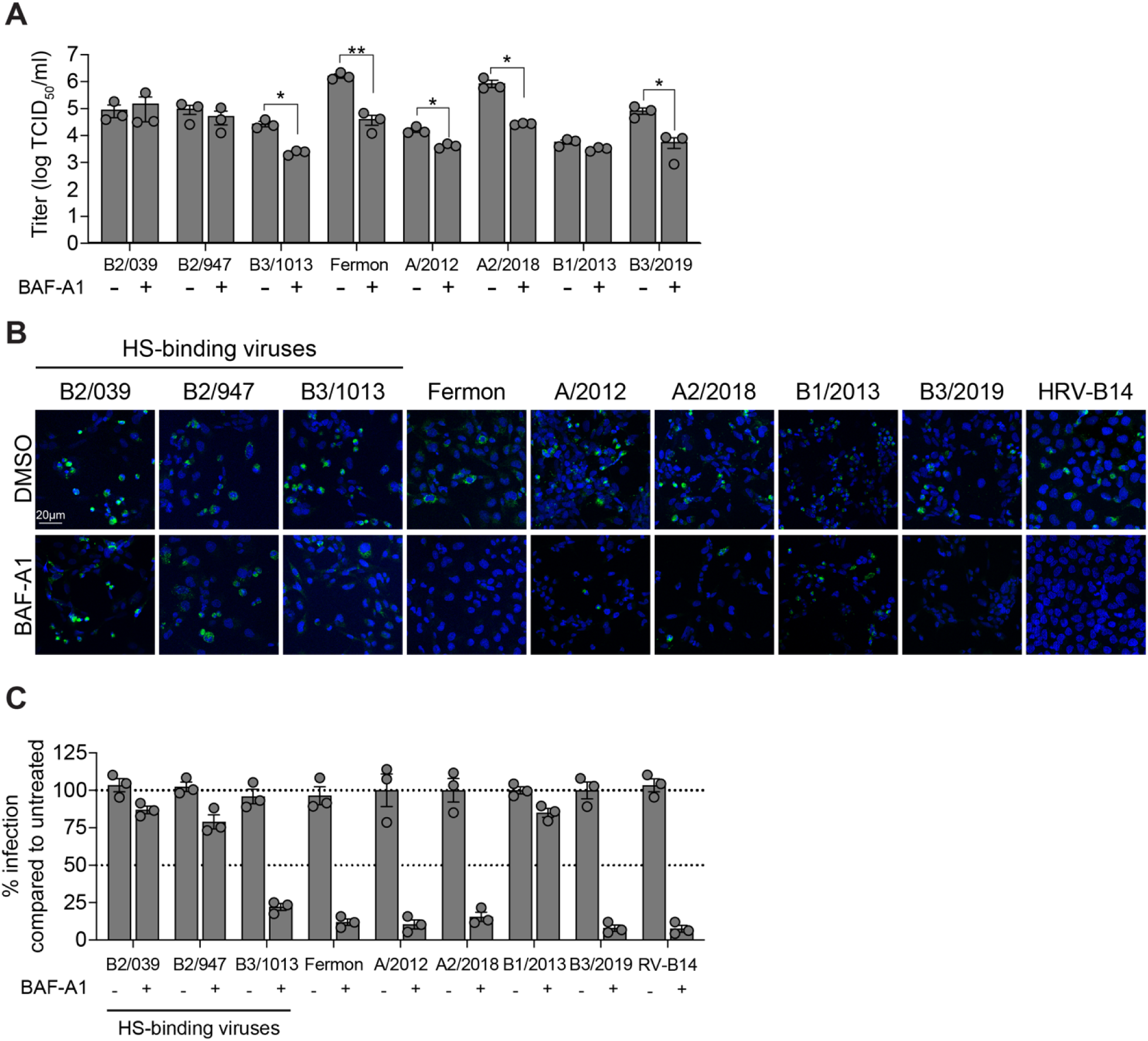
Acid sensitivity of Enterovirus-D68 strains. (A) RD cells were pretreated with 200□nM bafilomycin A1 (BAF-A1) for 1hour at 37°C followed by infection with Enterovirus-D68 (EV-D68) strains at multiplicity of infection (MOI) 1. After 24 hrs, virus titers in cell lysates were determined by endpoint dilution. Data displayed represents averages of technical replicates ± standard error of the mean (SEM) and are derived from three independent experiments. Statistical analysis was performed with Students t-test comparing untreated to BAF-A1 treated infected cells. P > 0.05; * P ≤ 0.05; ** P ≤ 0.01; *** P ≤ 0.001; **** P ≤ 0.0001 (B) RD cells were pretreated with 200 nM BAF-A1 for 1hour at 37°C followed by infection with EV-D68 strains at MOI 1. After 24hrs, cells were fixed and stained for the presence of double stranded RNA (green) and nuclei were visualized using Hoechst (blue). (C) Quantification of infection percentage comparing EV-D68 infected RD cells with untreated or BAF-A1-pretreatment. Data displayed represents averages of technical replicates ± standard error of the mean (SEM) and are derived from three independent experiments. BAF-A1, bafilomycin A1; EV-D68, Enterovirus-D68; MOI, multiplicity of infection; SEM, standard error of mean.

## Discussion

Here, we examine the receptor specificity of EV-D68 strains from various clades towards HS, N-glycans and glycolipid structures at the cell membrane. These data confirmed that only EV-D68 strains that carry an amino acid substation in VP1 at position 271 that renders a negative charge into a positive one can bind to HS. Our research revealed that EV-D68 strains bind not only to α2,6-linked N-glycans, but also to glycan structures terminating in α2,8-linked sialosides, such as the disialylated gangliosides GD3, GD1c, GT1a, and GQ1b, which serve as additional receptors.

EV-D68 binding to HS has been considered a cell culture adaptation associated with an amino acid substitution E271K in VP1^26,27^. HS carries a net negative charge and the previously reported EV-D68 strain B2/947 and B3/1013 have a positively charged lysine in VP1 position 271 instead of the negatively charged glutamic acid^26,27^. Here, we identified another amino acid substitution in VP1 E271R rendering glutamic acid into the positively charged arginine in the HS-binding EV-D68 strain B2/039. This indicates that EV-D68 strains can have different amino acid polymorphism at position 271 that facilitate HS binding, but it seems to be restricted to positively charged residues. The relevance of HS-binding for the pathogenesis of EV-D68 is unclear. Phenotypically the HS-binding of EV-D68 resulted in increased replication efficiency in neuroblastoma and RD cell lines^27^, but did not show an replication advantage in more physiological models such as human primary airway and brain organoids^48^. No amino acid polymorphism at position 271 in VP1 is observed in the Nextstrain database^40^ (Supplementary Figure 5), which raises the possibility that HS binding is not a relevant characteristic in circulating strains. However, initially believed that HS binding in EV-A71 is only a cell culture adaptation^49^, it was shown that it emerged in an immunocompromised host, and affects the pathogenesis^50^. Whether this is the case for EV-D68 remains to be established EV-D68 has previously been shown to bind α2,3-, and α2,6-linked SIAs^23–25^. Our SIA glycan array consisted of α2,3-, α2,6-linked SIAs and ganglioside structures that contain both α2,3- and α2,8-linked SIA. Several EV-D68 strains showed high fluorescence intensity for specifically the disialylated gangliosides GD3, GD1c, GT1a, GQ1b. Importantly, the lack of α2,3-linked sialic acid binding in our studies is consistent with previously published results^23,24^. The trait of EV-D68 to bind to SIA is not unique among the species *Enterovirus deconjuncti.* It has been shown that EV-D70 recognizes α2,3-linked SIA^51^ while EV-D94 recognizes α2,3- and α2,6-linked SIA^23^. Which sialyltransferases are important remain to be established, but it is a possibility that ST3 sialyltransferases are used as these create the acceptors for ST8 sialyltransferases installing the α2,8-linked SIA that we identified to be receptors for enteroviruses. α2,8-linked SIA is predominantly found on gangliosides and are used other viruses within the family *Picornaviridae* such as porcine sapelovirus^52^ and hepatitis A virus^53^. Coronaviruses and influenza A viruses have also been shown to functionally interact with gangliosides^54–60^. This is the first report of an enterovirus that uses disialylated gangliosides as an additional receptor. So far, we consider that ganglioside binding is not related to cell culture adaptation, as also EV-D68 strains without affinity to HS bind to gangliosides. The exact role of gangliosides in viral entry is not yet understood, but we speculate that gangliosides induce uncoating of EV-D68 strains potentially using a similar mechanism as hepatitis A virus.

EV-D68 primarily infects cells of the human respiratory tract and is mainly associated with mild disease. In certain cases severe respiratory disease as well as neurological complications such as encephalitis, meningitis, acute flaccid paralysis and rarely Guillain-Barré can occur^61,62^. EV-D68 strains are able to infect and spread in motor neurons, which is thought to be independent of SIA as removal of SIA with NA did not affect virus growth^63,64^. By continuous endocytosis gangliosides get internalized and can be found in endosomes which might facilitate that gangliosides can support uncoating of viruses within neurons even after NA treatment. The potential role of ganglioside as uncoating receptors needs to be established though.

Taken together, we provide strong evidence that HS recognition is acquired during cell culture passaging as a positive charge at position 271 of VP1 is identified in all virus stocks with HS binding properties. To a certain extent the virus strains also show differences their phenotypic characteristics Thus, we highly recommend sequencing all virus stocks before usage in experimental studies. Furthermore, we show that EV-D68 uses a wide range of glycan structures and newly glyolipids as entry receptors. This highlights that there is a large promiscuity in the recognition of glycan structures among different EV-D68 strains. How and whether this variation affects the pathogenesis of EV-D68 is an important next question to address.

## Material and Methods

### Cells

HeLa-Rh cells (kindly provided by Johan Neyts from the Rega Institute at the KU Leuven) and Rhabdomyosarcoma (RD) cells (American Type Culture Collection (ATCC)) were maintained in Dulbecco’s Modified Eagle’s medium (DMEM; Lonza, Basel, Switzerland) supplemented with 10% dialysed fetal bovine serum (FBS, Sigma-Aldrich, St. Louis, MO, USA), 2mM L-glutamine (Lonza) and 1% penicillin/streptomycin (Lonza) at 37°C with 5% CO_2_. SK-N-SH cells (ATCC) and A549 (ATCC) were maintained in Eagle’s minimum essential medium (EMEM, Lonza) supplemented with 10% FCS (Lonza), 1% penicillin/streptomycin (Lonza), 2mM L-glutamine (Lonza), 1% nonessential amino acids (Lonza), 1mM sodium pyruvate (Thermo Fisher Scientific, Waltham, MA, USA) and 1,5 mg/mL sodium bicarbonate (Lonza) at 37°C with 5% CO_2_. Medium was refreshed every 2–4 days, and cells were passaged at >80% confluence using PBS and trypsin-EDTA. Cells were regularly checked for presence of mycoplasma.

### Reagents

GENZ-123346, Bafilomycin A1 and (*S*)-fluoxetine were purchased from Sigma Aldrich and dissolved in dimethylsulfoxide (DMSO, Sigma Aldrich) at a concentration of 10 mM. Heparan (Sigma-Aldrich, H4784) was dissolved in H_2_O at a concentration of 30 mg/mL.

### Viruses

Enterovirus D68 (EV-D68) strains were derived from patients diagnosed with EV-D68 infection at the National Institute of Public Health and the Environment (RIVM, Bilthoven, The Netherlands). For *in vitro* studies, virus stocks with accession numbers from Table 2 were grown in RD cells (ATCC) at 33°C in 5% CO_2_. The prototype EV-D68 strain Fermon strain was provided by Frank van Kuppeveld (Utrecht University, Utrecht, The Netherlands) and also propagated in RD cells. All EV-D68 strains utilized in this study originated from the same or a second laboratory passage. After passaging, the full genome was Sanger sequenced to check for cell culture adaptations. Infectious clone of human rhinovirus B14 (HRV-B14) (pWR3.26) was purchased from ATCC. HRV-B14 was obtained by linearizing the plasmid with MluI, following *in vitro* transcription of viral RNA using the T7 RiboMAX™ Express Large Scale RNA Production System according to the manufacturer’s protocol. To obtain infectious virus, *in vitro* transcribed RNA was transfected into HeLa-Rh cells using Lipofectamin 2000 (Invitrogen).

**Table 2:**
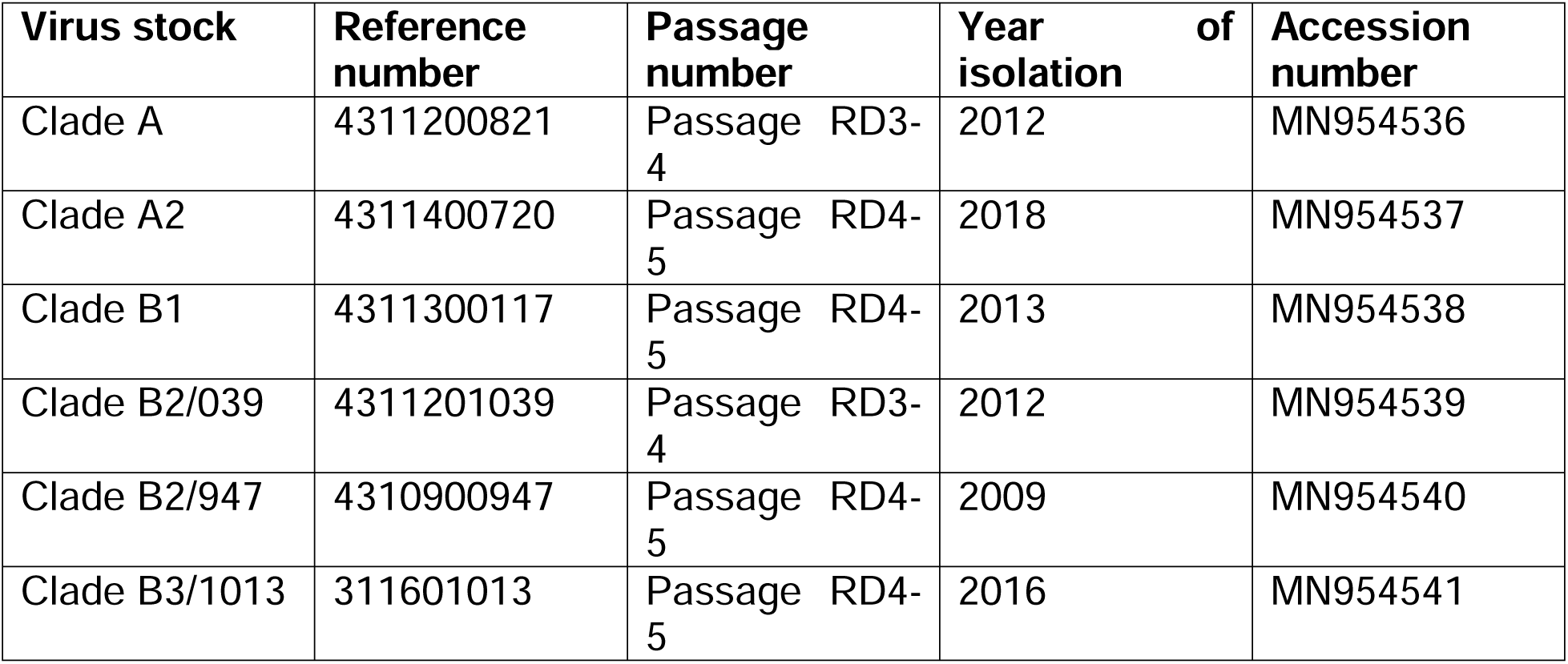
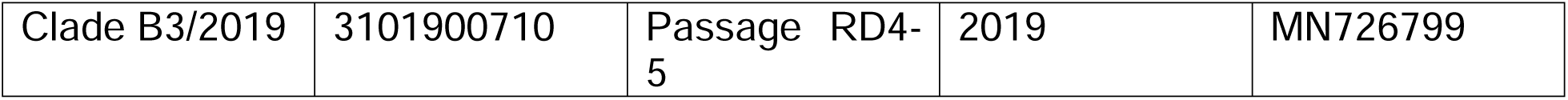
Virus strains, including reference number, passage number, year of isolation and accession number.

### Sanger sequencing of virus stock to detect cell culture adaptations

Following the manufacturer’s protocol, viral RNA was extracted using the Roche High Pure RNA Isolation Kit (Roche, The Netherlands). Subsequently, cDNA synthesis was performed using the Superscript IV First Strand Synthesis kit (Thermo Fischer Scientific) according to the manufacturer’s instructions. The synthesised cDNA was then subjected to AmpliTaq PCR to amplify the genome of the EV-D68 strains using the primer sets listed in Table 3. The initial denaturation step was conducted at 95°C for 6 minutes, followed by 40 cycles of denaturation at 45°C for 30 seconds, annealing at 55°C for 1 minute, and extension at 72°C for 2 minutes. A final extension at 72°C for 10 minutes was performed, and the products were stored at 4°C until further use. The PCR products were separated and analyzed on a 2% agarose gel, and the corresponding products were purified using the MinElute Gel Extraction Kit (Qiagen, Germantown, Maryland, USA) according to the manufacturer’s instructions. Subsequently, AbiSeq PCR was performed with 2µM of forward and reverse primers, Seq 3.1 Buffer, Big Dye, 2.5 µl sample, and bidest to adjust the final volume to 10 µl. The PCR involved 30 cycles of denaturation at 96°C for 10 seconds, annealing at 45°C for 30 seconds, and extension at 60°C for 4 minutes. Post-PCR, the sequence products were purified using a Sephadex plate (BD Biosciences) according to the manufacturer’s protocol. The purified products were then sequenced on the ABI3130XL sequencer. Sequence alignment was performed using the software program Unipro UGENE v.42.039.

**Table 3:**
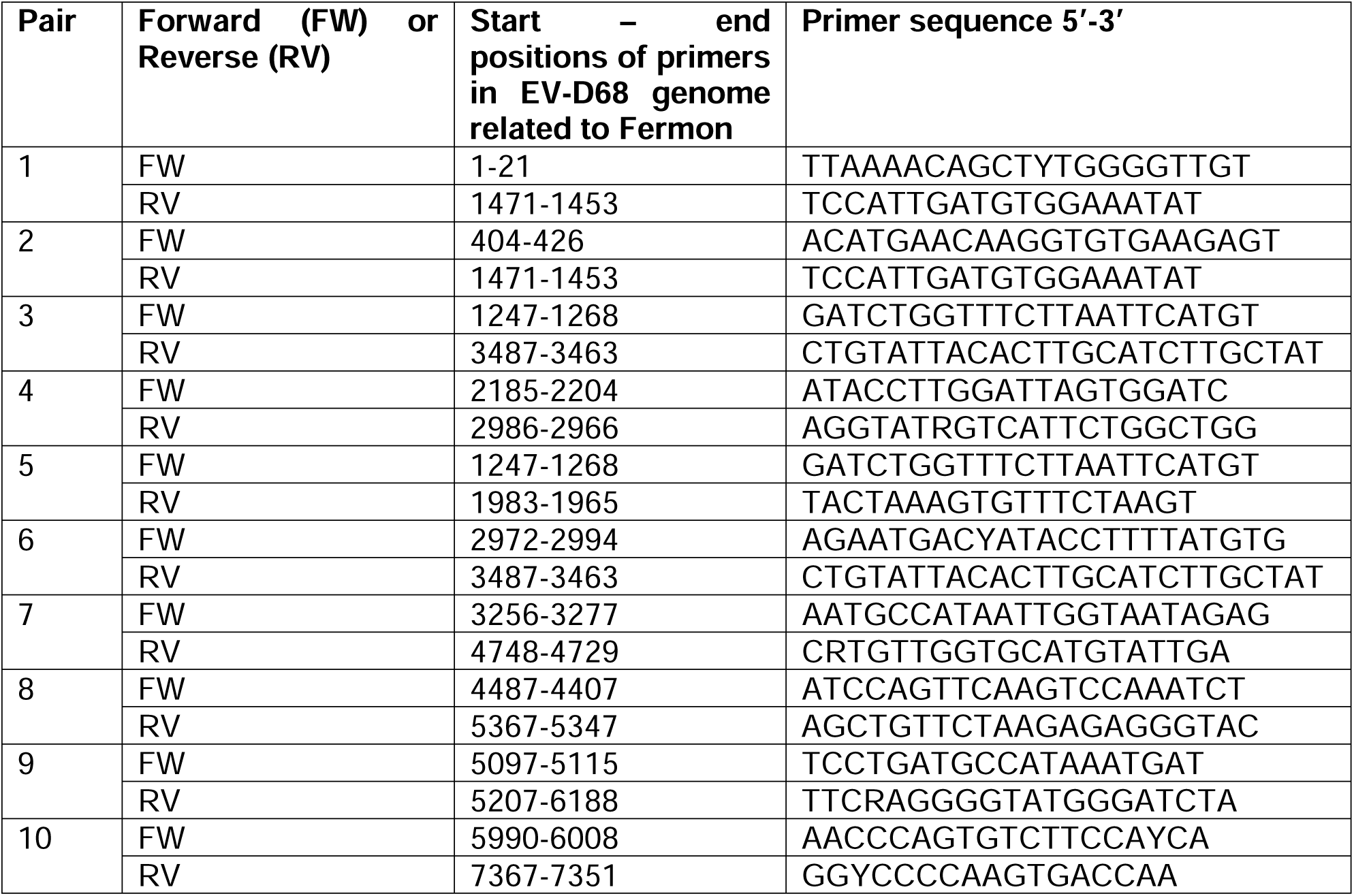
Primer sets spanning the genome of EV-D68.

### Glycan array analyses

For the glycan microarray studies, the printed library of compounds comprised the glycans and quality control procedures were published previously^41,65^. Virus strains (25 μL) were diluted with PBS-T (PBS + 0.1% Tween, 25 μL) and applied to the array surface in a humidified chamber for 1 h, followed by successive rinsing with PBS-T (PBS + 0.1% Tween), PBS and deionized water (2x) and dried by centrifugation. The virus-bound slide was incubated for 1 h with the polyclonal rabbit anti-Enterovirus D68 VP1 antibody (GeneTex, GTX132313, Lot Nr: 42284, 100 μL, 5 μg/mL in PBS-T). A secondary goat anti-rabbit AlexaFluor-647 antibody (100 μL, 2 μg/mL in PBS-T) (Thermo Fisher) was applied, incubated for 1 h in a humidified chamber and washed again as described above. Slides were dried by centrifugation after the washing step and scanned immediately using an Innopsys Innoscan 710 microarray scanner at the appropriate excitation wavelength. To ensure that all signals were in the linear range of the scanner’s detector and to avoid any saturation of the signals various gains and PMT values were employed. Images were analyzed with Mapix software (version 8.1.0 Innopsys) and processed with our home-written Excel macro. The average fluorescence intensity and SD were measured for each compound after exclusion of the highest and lowest intensities from the spot replicates (n = 4).

### Synthesis of multivalent ligands as inhibitors

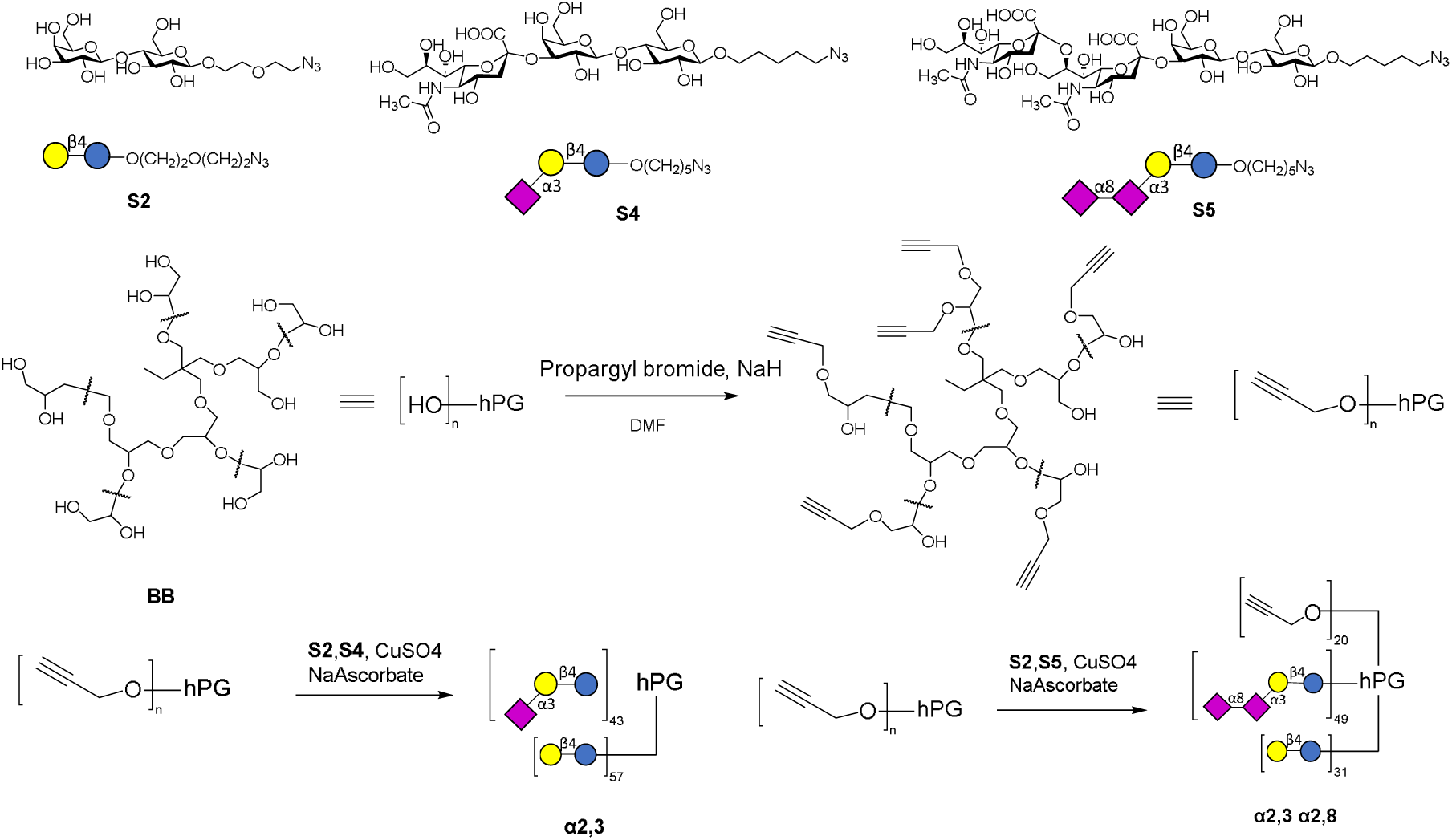

GM3 and GD3 oligosaccharides were chemoenzymatically synthesized from peracetylated lactose. Trichloroacetimidate was installed upon anomeric deprotection with hydrazine acetate. Subsequent TMSOTf promoted glycosylation, followed by zemplen deacetylation yielded lactose functionalized with two different azido linkers (**S2**) and (**S3**). Lactose functionalized with an azidopentanol linker (**S3**) was elongated to GM3 (**S4**) and GD3 (**S5**) based oligosaccharides using PmST1 M144D and CST-II as described before^66^.

Hyperbranched polyglycerol (hPG) was synthesized from tris(hydroxymethyl)propane through anionic polymerization. Careful use of solvent and temperature yielded a hPG with Mn of 15kDa as the backbone (**BB**)^67^, determined though ^1^H NMR and inverse gate ^13^C NMR as described before^67,68^. Introduction of propargyl groups on the barckbone yielded a propargyl functionalized hPG with a functionalization degree of 60% as determined by ^1^H NMR. Azido functionalized glycans were conjugated to alkyne-hPG using CuAAC. Composition of the polymeric glycans was determined through comparing characteristic protons in ^1^H NMR to yield glycopolymers functionalized with GM3 (hPG-α**2,3**) and GD3 (hPG-α**2,3-**α**2,8**).

### Single Cycle Virus Infection

Virus infections were performed by incubating the corresponding cells with virus at the indicated MOI at 33 °C for 60 minutes. After incubation, the inoculum was removed, and cells were washed three times with PBS. At the indicated time points either supernatants were collected or cells were frozen. For measurements of infectious particles of cell lysates, virus was released from the cells by three freeze-thawing cycles.

### Virus Titrations

Virus titers of virus stocks, supernatant or cell lysates were determined by endpoint dilution. Briefly, 10-fold dilutions were prepared on a subconfluent layer of RD cells and cells were incubated at 33°C in 5% CO2. Cytopathic effect (CPE) was was visually inspected and virus titers were determined and calculated by the method of Reed and Muench and expressed as 50% tissue culture infectious dose (TCID50)^69^.

### Removal of SIAs, glycolipids and Heparan Sulfate

RD, SK-N-SH and A549 cells were incubated with 100mU/mL *Arthrobacter ureafaciens* neuraminidase (Roche) in FBS-free cell culture medium at 37°C in 5% CO_2_ for 1 hour to remove SIAs. To specifically deplete gangliosides and α2,8-linked SIAs (SIAs) on glycoproteins, RD, SK-N-SH and A549 cells were treated for 72 hours at 37°C in a 5% CO_2_ with 5 µM GENZ-123346. To remove heparan sulfate and prevent sulfation, RD, SK-N-SH and A549 cells were incubated with 80mM NaClO3 (Sigma Aldrich) for at least two weeks at 37°C in a 5% CO_2_ incubator.

### Flow Cytometry detection of ganglioside expression

The reduction of glyoclipids was confirmed using flow cytometry (FACS). Following trypsinisation, cells were washed with 100µl of FACS buffer (PBS with MgCl2 and CaCl2, 2mM EDTA, and 0.05% BSA). After washing, cells were incubated with FACS Buffer II containing PBS with MgCl2 and CaCl2, 2mM EDTA, and 0.1% BSA and 10µg/mL CTX-FITC (C1655, Sigma-Aldrich). The cells were incubated for 30 minutes at 4°C and protected from light. Following incubation, the cells were centrifuged at 1250 rpm for 5 minutes at 4°C. The supernatant was discarded, and the cells were washed again with 100 µl of FACS Buffer II, followed by centrifugation and removal of the supernatant. Finally, cells were resuspended in 150µl of FACS buffer and analysed using the FACSlyric Flow Cytometer (BD Biosciences, Franklin Lakes, New Jersey, USA).

### Heparin-binding assay

EV-D68 strains were either incubated with heparin(1mg/mL) or plain medium for 1 h at 37°C. RD cells were infected with Heparin-treated (1mg/mL) and untreated viruses at MOI 1. After 1 h of incubation at 33°C, inoculum was taken off and cells were washed three times with PBS. 24 hours post-inoculation cells were frozen and virus titer of cell lysates was determined by endpoint dilution. Additionally, cells were fixed 24 hrs post-infection and subjected to immunofluorescence stainings.

### Inhibition assay

EV-D68 strains were incubated with the glycolipid mimetics GM3 and GD3 or the corresponding empty backbone structure or lactose for 1h at 37°C. After the incubation RD cells were infected with viruses at MO1. Virus titer of cell lysates was determined by endpoint dilution 24 hrs post infection.

### BAF-A1 treatment assay

Prior to infection, RD cells were treated with 200nM Bafilomycin A1 (BAF-A1) or DMSO for 1h to inhibit vascular acidification. Afterwards, RD cells were infected with EV-D68 viruses at MOI1. After 1hour of infection, virus inoculum was removed and cells were washed three times with PBS. After washing, DMEM containing 200nM BAF-A1 or DMSO was added. 24 hours post-infection, the cells on glass slides were fixed and supernatant was collected. The glass slides were further processed for immunofluorescent staining, and infectious titer in the supernatant was determined by endpoint dilution.

### Immunofluorescence Microscopy

RD cells on glass slides were fixed with 10% formalin for 30 minutes at room temperature. After fixation, the cells were washed with PBS and then permeabilised with PBS containing 1% Triton X-100 (Sigma Aldrich). Following permeabilisation, the cells were blocked for 30 minutes at RT with a Washing Buffer consisting of PBS, 0.5% Triton X-100 and 1% BSA (Sigma Aldrich). The primary antibodies for double stranded RNA (Sigma Aldrich, clone rJ2, MABE1134) was used at a concentration of 2,5 µg/mL and was diluted in this Washing Buffer and incubated at RT for 1h. Subsequently, the glass slides were washed three times in PBS before staining with 0,2 µg/mL of the secondary antibodies donkey anti-Mouse IgG (H+L) Highly Cross-Adsorbed Secondary Antibody, Alexa Fluor™ 488 (A-21202, Invitrogen) and 2µM Hoechst (Life Technologies/Invitrogen, H3570) dye, diluted in Washing Buffer, for 1h at room temperature. Lastly, the stained cells were washed three times in PBS and once in water, mounted in ProLong Antifade Mountant, and imaged using a Zeiss LSM 700 confocal microscope.

### Confocal image quantification

To quantify the number of infected cells, the confocal pictures were blinded. For each sample, three images were captured, and each experiment was performed three independent times in technical triplicate. Afterwards, the pictures were blinded by one of the authors and counted by another author. Infection percentage was determined by counting nuclei (Hoechst) and infected cells (double stranded RNA staining).

### Temperature Sensitivity assay

To assess the thermostability of EV-D69 strains, viruses were subjected to a temperature gradient ranging from 35°C to 60°C for 15 minutes. To assess HS binding, viruses were treated with 1mg/mL heparin and subjected to the same temperature gradient. After temperature exposure, EV-D68 infectivity was assessed by endpoint dilution.

### pH Sensitivity assay

Virus stocks were subjected to a pH gradient ranging from pH 4 to pH 7.4 to determine the acid sensitivity. pH was adjusted carefully using sterile hydrochloric acid (HCl; 1M). The accuracy of the pH was confirmed with pH test strips (VWR Chemicals). Where mentioned, viruses were treated with 1mg/mL heparin and subjected to pH titration. After acidification, 20 µL of the viral stocks were titrated in 200 µL of DMEM, which neutralised the pH without further adjustments. EV-D68 infectivity was assessed by endpoint dilution

### Sequence Alignments

The sequences were derived from the Sanger sequencing reads generated from the virus stocks. The protein sequences were manually annotated and VP1 and VP3 amino acid sequences were extracted. Sequences were aligned with Clustal OMEGA^70^ and sequences similarities and secondary structure information were analysed with ESPRIPT3.0^71^ and Chimera^72^

### Calculations and Statistics

Each experiment was performed in technical triplicates, with at least three independent experiments. Statistical significance was determined using one-way ANOVA, paired t-test, or unpaired t-test as indicated with a threshold of P < 0.05 considered significant. Statistical analyses, nonlinear regression, dose-response inhibition, and graphical representations were performed using GraphPad Prism Version 10.2.3 (La Jolla, CA, USA) and Adobe Illustrator. Chimera was used to visualize the protein structures PDB: 5BNP and 5BNP.

## Supporting information

Supplement Part

## Acknowledgement

We thank Kristina Lanko and Ruben Hulswit for technical assistance, scientific discussions and for critically reading our manuscript. L.B is supported by a fellowship from The Netherlands Organization for Scientific Research (VENI contract 09150162210154). D.v.R is supported by fellowships from the Netherlands Organization for Scientific Research (VIDI contract 91718308).

## Contributions

Conceptualization RPdV, DvR, LB

Investigation ASKPdA, JvT, AM, SSNA, LB

Formal Analysis ASKPdA, JvT, AM, SSNA, LB

Resources GJB, DvR, RPdV, LB

Methodology ASKPdA, JvT, AM, JF, SSNA, RPdV, LB

Supervision RPdV, RJP, GJB, LB

Visualization ASKPdA, JvT, RPdV, LB

Writing Original ASKPdA, JvT, RPdV, LB

Writing-Reviewing all authors

Funding acquisition RPdV, LB

## References

1. Grizer, C. S., Messacar, K. & Mattapallil, J. J. Enterovirus-D68 – a reemerging non-polio enterovirus that causes severe respiratory and neurological disease in children. Front. Virol. 4, (2024).

2. Sooksawasdi Na Ayudhya, S., Laksono, B. M. & van Riel, D. The pathogenesis and virulence of enterovirus-D68 infection. Virulence 12, 2060–2072 (2021).

3. Cassidy, H., Poelman, R., Knoester, M., Van Leer-Buter, C. C. & Niesters, H. G. M. Enterovirus D68 – The New Polio? Frontiers in Microbiology 9, (2018).

4. Khetsuriani, N., Lamonte-Fowlkes, A., Oberst, S., Pallansch, M. A., & Centers for Disease Control and Prevention. Enterovirus surveillance--United States, 1970-2005. MMWR Surveill Summ 55, 1–20 (2006).

5. Itagaki, T. et al. Seroprevalence of enterovirus D68 in Yamagata, Japan, between 1976 and 2019. J Med Virol 96, e29947 (2024).

6. Matthew R. Vogt; Peter Wright; William Hickey; James E. Crowe, J. K. B. Enterovirus D68 RNA Visualized in the Anterior Horn of the Spinal Cord of a Pediatric Patient with Flaccid Paralysis. 7, 712 (2020).

7. Midgley, C. M. et al. Severe respiratory illness associated with a nationwide outbreak of enterovirus D68 in the USA (2014): a descriptive epidemiological investigation. The Lancet Respiratory Medicine 3, 879–887 (2015).

8. Edwin, J. J. et al. Surveillance summary of hospitalized pediatric enterovirus D68 cases in Canada, September 2014. Can Commun Dis Rep 41, 2–8 (2015).

9. Holm-Hansen, C. C., Midgley, S. E. & Fischer, T. K. Global emergence of enterovirus D68: a systematic review. The Lancet Infectious Diseases 16, e64–e75 (2016).

10. Xiao, Q. et al. Prevalence and molecular characterizations of enterovirus D68 among children with acute respiratory infection in China between 2012 and 2014. Sci Rep 5, 16639 (2015).

11. Thongpan, I. et al. Prevalence and Phylogenetic Characterization of Enterovirus D68 in Pediatric Patients with Acute Respiratory Tract Infection in Thailand. Jpn J Infect Dis 69, 426–430 (2016).

12. Huang, Y.-P., Lin, T.-L., Lin, T.-H. & Wu, H.-S. Molecular and epidemiological study of enterovirus D68 in Taiwan. Journal of Microbiology, Immunology and Infection 50, 411–417 (2017).

13. Benschop, K. S. et al. Re-emergence of enterovirus D68 in Europe after easing the COVID-19 lockdown, September 2021. Eurosurveillance 26, 2100998 (2021).

14. Jallow, M. M. et al. Real-Time Enterovirus D68 Outbreak Detection through Hospital Surveillance of Severe Acute Respiratory Infection, Senegal, 2023. Emerg Infect Dis 30, 1687–1691 (2024).

15. Xie, Z., Khamrin, P., Maneekarn, N. & Kumthip, K. Epidemiology of Enterovirus Genotypes in Association with Human Diseases. Viruses 16, 1165 (2024).

16. Shah, M. M. Enterovirus D68-Associated Acute Respiratory Illness ─ New Vaccine Surveillance Network, United States, July–November 2018–2020. MMWR Morb Mortal Wkly Rep 70, (2021).

17. Cao, R. G., Mejias, A., Leber, A. L. & Wang, H. Clinical and molecular characteristics of the 2022 Enterovirus-D68 outbreak among hospitalized children, Ohio, USA. J Clin Virol 169, 105618 (2023).

18. Fall, A. et al. Enterovirus D68: Genomic and Clinical Comparison of 2 Seasons of Increased Viral Circulation and Discrepant Incidence of Acute Flaccid Myelitis— Maryland, USA. Open Forum Infectious Diseases 11, ofae656 (2024).

19. Simoes, M. P. et al. Epidemiological and Clinical Insights into the Enterovirus D68 Upsurge in Europe 2021–2022 and Emergence of Novel B3-Derived Lineages, ENPEN Multicentre Study. The Journal of Infectious Diseases 230, e917–e928 (2024).

20. Hodcroft, E. B. et al. Evolution, geographic spreading, and demographic distribution of Enterovirus D68. PLoS Pathog 18, e1010515 (2022).

21. Wei, W. et al. ICAM-5/Telencephalin Is a Functional Entry Receptor for Enterovirus D68. Cell Host and Microbe 20, 631–641 (2016).

22. Liu, X. et al. MFSD6 is an entry receptor for respiratory enterovirus D68. Cell Host & Microbe (2025) doi:10.1016/j.chom.2024.12.015.

23. Baggen, J. et al. Enterovirus D68 receptor requirements unveiled by haploid genetics. Proceedings of the National Academy of Sciences of the United States of America 113, 1399–1404 (2016).

24. Imamura, T. et al. Antigenic and Receptor Binding Properties of Enterovirus 68. Journal of Virology 88, 2374–2384 (2014).

25. Liu, Y. et al. Sialic acid-dependent cell entry of human enterovirus D68. Nat Commun 6, 8865 (2015).

26. Baggen, J. et al. Bypassing pan-enterovirus host factor PLA2G16. Nature Communications 2019 10:1 10, 1–10 (2019).

27. Sooksawasdi Na Ayudhya, S., et al. Enhanced Enterovirus D68 Replication in Neuroblastoma Cells Is Associated with a Cell Culture-Adaptive Amino Acid Substitution in VP1. mSphere 5, e00941–20 (2020).

28. Elrick, M. J., Pekosz, A. & Duggal, P. Enterovirus D68 molecular and cellular biology and pathogenesis. Journal of Biological Chemistry 296, 100317 (2021).

29. Gahmberg, C. G., Tian, L., Ning, L. & Nyman-Huttunen, H. ICAM-5--a novel two-facetted adhesion molecule in the mammalian brain. Immunol Lett 117, 131–135 (2008).

30. Bagchi, S. et al. Probable role for major facilitator superfamily domain containing 6 (MFSD6) in the brain during variable energy consumption. International Journal of Neuroscience 130, 476–489 (2020).

31. Harduin-Lepers, A. The vertebrate sialylation machinery: structure-function and molecular evolution of GT-29 sialyltransferases. Glycoconj J 40, 473–492 (2023).

32. Krzewinski-Recchi, M.-A. et al. Identification and functional expression of a second human β-galactoside α2,6-sialyltransferase, ST6Gal II. European Journal of Biochemistry 270, 950–961 (2003).

33. Mohamed, K. A., Kruf, S. & Büll, C. Putting a cap on the glycome: Dissecting human sialyltransferase functions. Carbohydr Res 544, 109242 (2024).

34. Sipione, S., Monyror, J., Galleguillos, D., Steinberg, N. & Kadam, V. Gangliosides in the Brain: Physiology, Pathophysiology and Therapeutic Applications. Front. Neurosci. 14, (2020).

35. Schjoldager, K. T., Narimatsu, Y., Joshi, H. J. & Clausen, H. Global view of human protein glycosylation pathways and functions. Nat Rev Mol Cell Biol 21, 729–749 (2020).

36. Smits, N. C., Shworak, N. W., Dekhuijzen, P. N. R. & van Kuppevelt, T. H. Heparan Sulfates in the Lung: Structure, Diversity, and Role in Pulmonary Emphysema. The Anatomical Record 293, 955–967 (2010).

37. Condomitti, G. & Wit, J. de. Heparan Sulfate Proteoglycans as Emerging Players in Synaptic Specificity. Frontiers in Molecular Neuroscience 11, 14 (2018).

38. Wang, X. et al. A sensor-adaptor mechanism for enterovirus uncoating from structures of EV71. Nat Struct Mol Biol 19, 424–429 (2012).

39. Prchla, E., Kuechler, E., Blaas, D. & Fuchs, R. Uncoating of human rhinovirus serotype 2 from late endosomes. J Virol 68, 3713–3723 (1994).

40. Nextstrain: real-time tracking of pathogen evolution | Bioinformatics | Oxford Academic. https://academic.oup.com/bioinformatics/article/34/23/4121/5001388?login=false.

41. Liu, L. et al. Heparan Sulfate Proteoglycans as Attachment Factor for SARS-CoV-2. ACS Central Science 7, 1009–1018 (2021).

42. Baeuerle, P. A. & Huttner, W. B. Chlorate — a potent inhibitor of protein sulfation in intact cells. Biochemical and Biophysical Research Communications 141, 870–877 (1986).

43. Bauer, L. et al. Fluoxetine Inhibits Enterovirus Replication by Targeting the Viral 2C Protein in a Stereospecific Manner. ACS Infect Dis 5, 1609–1623 (2019).

44. Hurdiss, D. L. et al. Fluoxetine targets an allosteric site in the enterovirus 2C AAA+ ATPase and stabilizes a ring-shaped hexameric complex. Science Advances 8, eabj7615 (2022).

45. Scheibner, D. et al. Phenotypic effects of mutations observed in the neuraminidase of human origin H5N1 influenza A viruses. PLOS Pathogens 19, e1011135 (2023).

46. Broszeit, F. et al. Glycan remodeled erythrocytes facilitate antigenic characterization of recent A/H3N2 influenza viruses. Nat Commun 12, 5449 (2021).

47. Tee, H. K. et al. Enterovirus A71 adaptation to heparan sulfate comes with capsid stability tradeoff. 2024.02.23.581741 Preprint at 10.1101/2024.02.23.581741 (2024).

48. Enterovirus D68 Infection in Human Primary Airway and Brain Organoids: No Additional Role for Heparan Sulfate Binding for Neurotropism. Microbiology Spectrum 10, (2022).

49. Tseligka, E. D. et al. A VP1 mutation acquired during an enterovirus 71 disseminated infection confers heparan sulfate binding ability and modulates ex vivo tropism. PLoS Pathog 14, e1007190 (2018).

50. Weng, K.-F. et al. Variant enterovirus A71 found in immune-suppressed patient binds to heparan sulfate and exhibits neurotropism in B-cell-depleted mice. Cell Reports 42, 112389 (2023).

51. Alexander, D. A. & Dimock, K. Sialic acid functions in enterovirus 70 binding and infection. J Virol 76, 11265–11272 (2002).

52. Kim, D.-S. et al. Porcine Sapelovirus Uses α2,3-Linked Sialic Acid on GD1a Ganglioside as a Receptor. Journal of Virology 90, 4067–4077 (2016).

53. Das, A. et al. Gangliosides are essential endosomal receptors for quasi-enveloped and naked hepatitis A virus. Nat Microbiol 5, 1069–1078 (2020).

54. Li, Z. et al. Synthetic O-acetylated sialosides facilitate functional receptor identification for human respiratory viruses. Nature Chemistry 2021 13:5 13, 496–503 (2021).

55. Pronker, M. F. et al. Sialoglycan binding triggers spike opening in a human coronavirus. Nature 624, 201–206 (2023).

56. Tomris, I. et al. The HCoV-HKU1 N-Terminal Domain Binds a Wide Range of 9-O-Acetylated Sialic Acids Presented on Different Glycan Cores. ACS Infect Dis 10, 3880–3890 (2024).

57. Matrosovich, M. et al. Gangliosides are not essential for influenza virus infection. Glycoconj J 23, 107–113 (2006).

58. Matrosovich, M. N. et al. Avian influenza A viruses differ from human viruses by recognition of sialyloligosaccharides and gangliosides and by a higher conservation of the HA receptor-binding site. Virology 233, 224–234 (1997).

59. Nguyen, L. et al. Sialic acid-containing glycolipids mediate binding and viral entry of SARS-CoV-2. Nature Chemical Biology 2021 18:1 18, 81–90 (2021).

60. Tomris, I. et al. SARS-CoV-2 Spike N-Terminal Domain Engages 9-O-Acetylated α2-8-Linked Sialic Acids. ACS Chem Biol 18, 1180–1191 (2023).

61. Helfferich, J. et al. Acute flaccid myelitis and Guillain–Barré syndrome in children: A comparative study with evaluation of diagnostic criteria. European Journal of Neurology 29, 593–604 (2022).

62. Williams, C. J. et al. Cluster of atypical adult Guillain-Barré syndrome temporally associated with neurological illness due to EV-D68 in children, South Wales, United Kingdom, October 2015 to January 2016. Euro Surveill 21, (2016).

63. Hixon, A. M., Clarke, P. & Tyler, K. L. Contemporary Circulating Enterovirus D68 Strains Infect and Undergo Retrograde Axonal Transport in Spinal Motor Neurons Independent of Sialic Acid. Journal of Virology (2019) doi:10.1128/JVI.00578-19.

64. Rosenfeld, A. B., Warren, A. L. & Racaniello, V. R. Neurotropism of enterovirus D68 isolates is independent of sialic acid and is not a recently acquired phenotype. mBio 10, (2019).

65. Arunkumar, G. A. et al. Functionality of the putative surface glycoproteins of the Wuhan spiny eel influenza virus. Nat Commun 12, 6161 (2021).

66. Chemoenzymatic Synthesis of 9NHAc-GD2 Antigen to Overcome the Hydrolytic Instability of O-Acetylated-GD2 for Anticancer Conjugate Vaccine Development - Wu - 2021 - Angewandte Chemie International Edition - Wiley Online Library. https://onlinelibrary.wiley.com/doi/abs/10.1002/anie.202108610.

67. Ul-Haq, M. I., Shenoi, R. A., Brooks, D. E. & Kizhakkedathu, J. N. Solvent-assisted anionic ring opening polymerization of glycidol: Toward medium and high molecular weight hyperbranched polyglycerols. Journal of Polymer Science Part A: Polymer Chemistry 51, 2614–2621 (2013).

68. Haksar, D. et al. Fighting Shigella by Blocking Its Disease-Causing Toxin. J. Med. Chem. 64, 6059–6069 (2021).

69. Reed, L. J. & Muench, H. A SIMPLE METHOD OF ESTIMATING FIFTY PER CENT ENDPOINTS12. American Journal of Epidemiology 27, 493–497 (1938).

70. Madeira, F. et al. The EMBL-EBI Job Dispatcher sequence analysis tools framework in 2024. Nucleic Acids Res 52, W521–W525 (2024).

71. Robert, X. & Gouet, P. Deciphering key features in protein structures with the new ENDscript server. Nucleic Acids Research 42, W320–W324 (2014).

72. Pettersen, E. F. et al. UCSF Chimera--a visualization system for exploratory research and analysis. J Comput Chem 25, 1605–1612 (2004).

