## Supplement Part for "Sialic acid-containing glycolipids extend the receptor repertoire of Enterovirus-D68"

**Pereirinha da Silva et al., 2025**

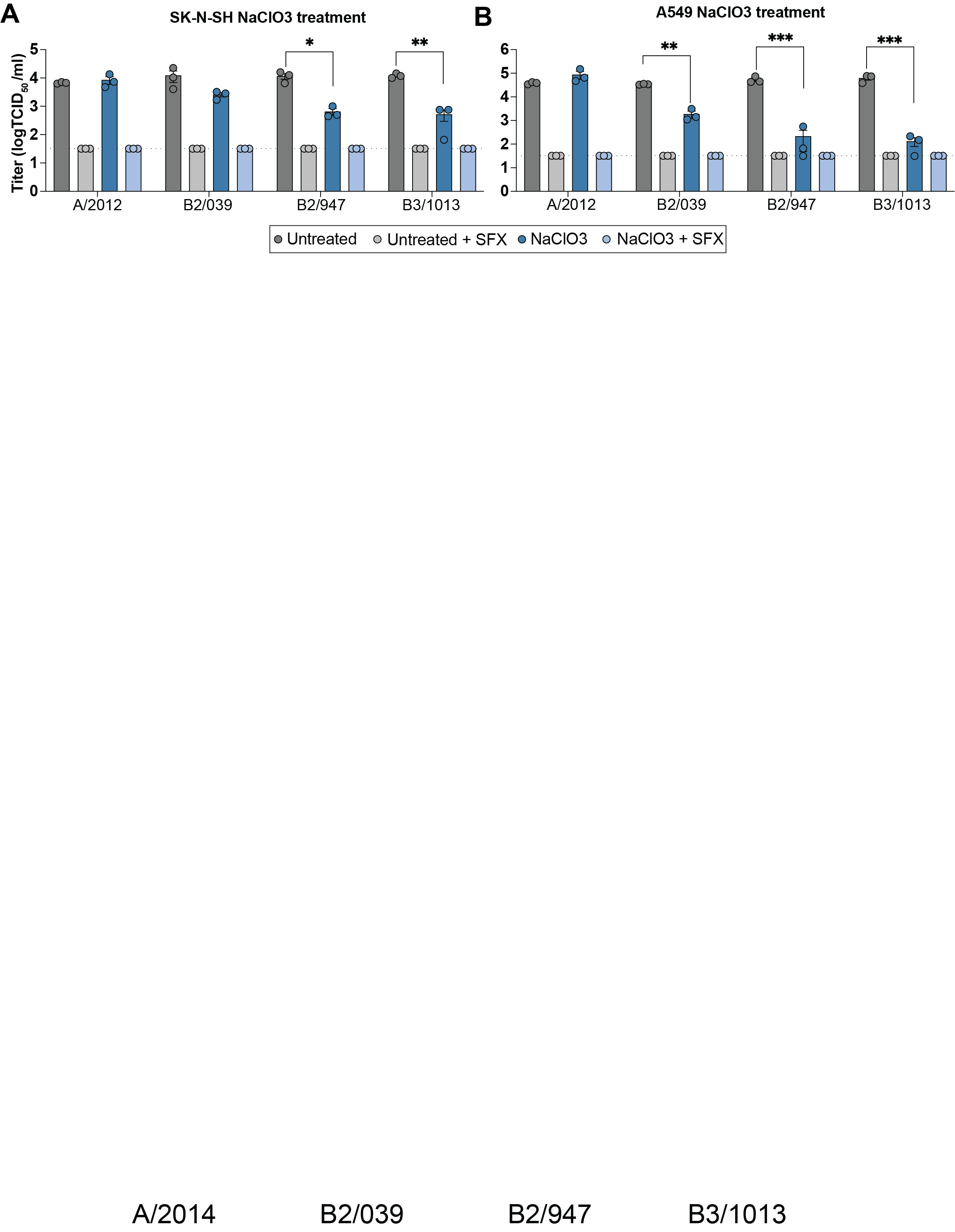

**Supplement Figure 1. Removal of Heparan Sulphate inhibits EV-D68 isolates B2/957 and B3/1013.** (A) SK-N-SH and (B) A549 cells were pre-treated and passaged in the presence of NaClO_3_ for at least two weeks. In a single-cycle viral replication assay, SK-N-SH and A549 cells were infected with Enterovirus-D68 isolates at multiplicity of infection 1. 10µM of the replication inhibitor (*S*)-fluoxetine (SFX) served as positive control. 24 hours post infection, cells were freeze-thawed three times, and virus titers in the lysates were determined by endpoint dilution. Data represent values from averaged technical replicates from three independent experiments ± the standard error of mean (SEM). Statistical analysis was performed with Students t-test comparing NaClO3 treated and infected cells to untreated cells upon infection. Asterisks indicate statistically significant differences P > 0.05; * P ≤ 0.05; ** P ≤ 0.01; *** P ≤ 0.001; **** P ≤ 0.0001

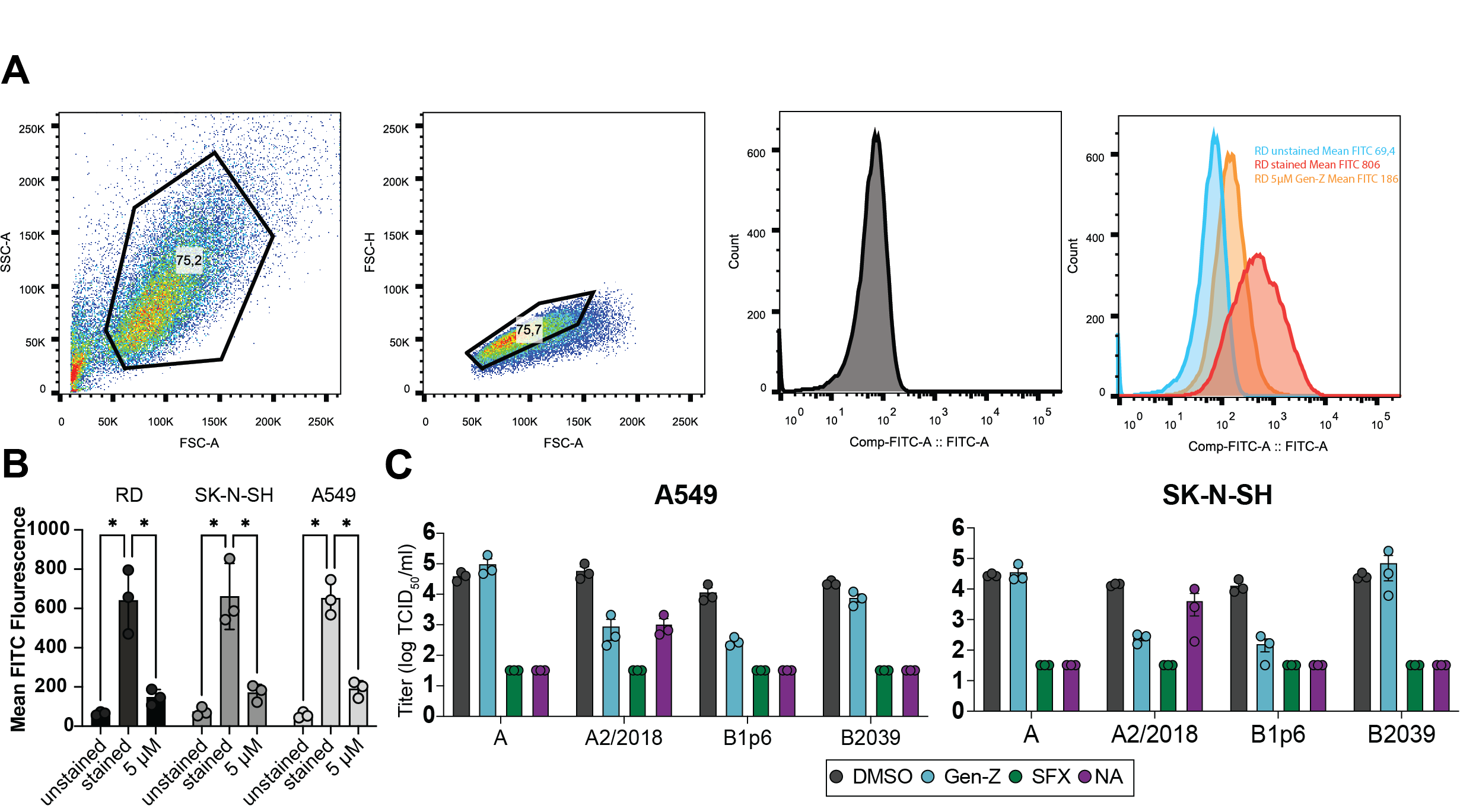

**Supplement Figure 2. Glycolipid depletion reduces replication of Enterovirus-D68 isolates that bind to gangliosides in different cell lines.** (A) RD, SK-N-SH and A549 cells were treated with 5µM Gen-Z123346 (Gen-Z) for 72 hours at 37°C. Glycolipids were detected by staining for the ganglioside GM-1 using FITC-labelled Cholera Toxin (CTX). Gating strategy to evaluate depletion of the ganglioside GM1. (B) Mean Fluorescent Values of CTX-FITC of RD, SK-N-SH and A459 cells after 72 hours of 5µM Gen-Z treatment. Data represented mean fluorescent values of three independent experiments. Statistical analysis was performed with One-Way ANOVA comparing cells to untreated stained control cells. (C) In a single-cycle viral replication assay, A549 and SK-N-SH cells were first pre-treated with 5µM Gen-Z123346 (Gen-Z) for 72hrs, 100 mU/ml *Arthrobacter ureafaciens* neuraminidase for 2hours. After pretreatment A549 and SK-N-SH cells were infected with Enterovirus-D68 isolates at multiplicity of infection 1. The replication inhibitor (*S*)-fluoxetine served as positive control. 24 hours post infection, virus titers of supernatants were determined by endpoint dilution. Data represent values from averaged technical replicates from three independent experiments ± SEM. Asterisks indicate statistically significant differences P > 0.05; * P ≤ 0.05; ** P ≤ 0.01; *** P ≤ 0.001; **** P ≤ 0.0001

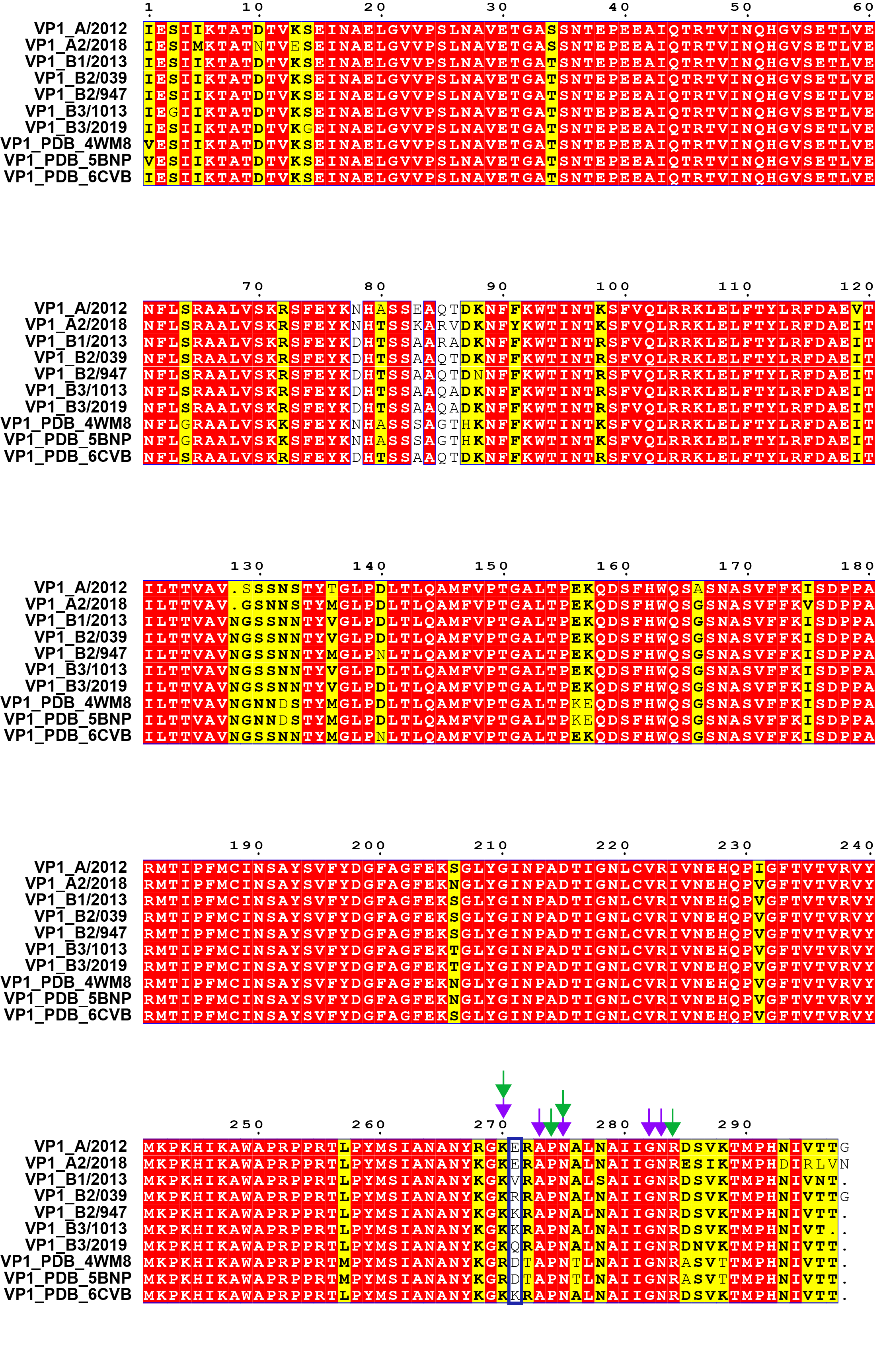

**Supplementary Figure 3. VP1 Multiple sequence alignment.** A multiple sequence alignment of the VP1 sequences of the sequenced Enterovirus-D68 virus stocks was performed with Clustal OMEGA^1^ . The ESPRIPT3.0 program^2^ was used to render sequence similarities. Purple (PDB: 5BNP) and green (PDB: 6CVB) arrows indicate residues in sialic acid coordination according to the Protein-Ligand Interaction Profiler PLIP^3^.

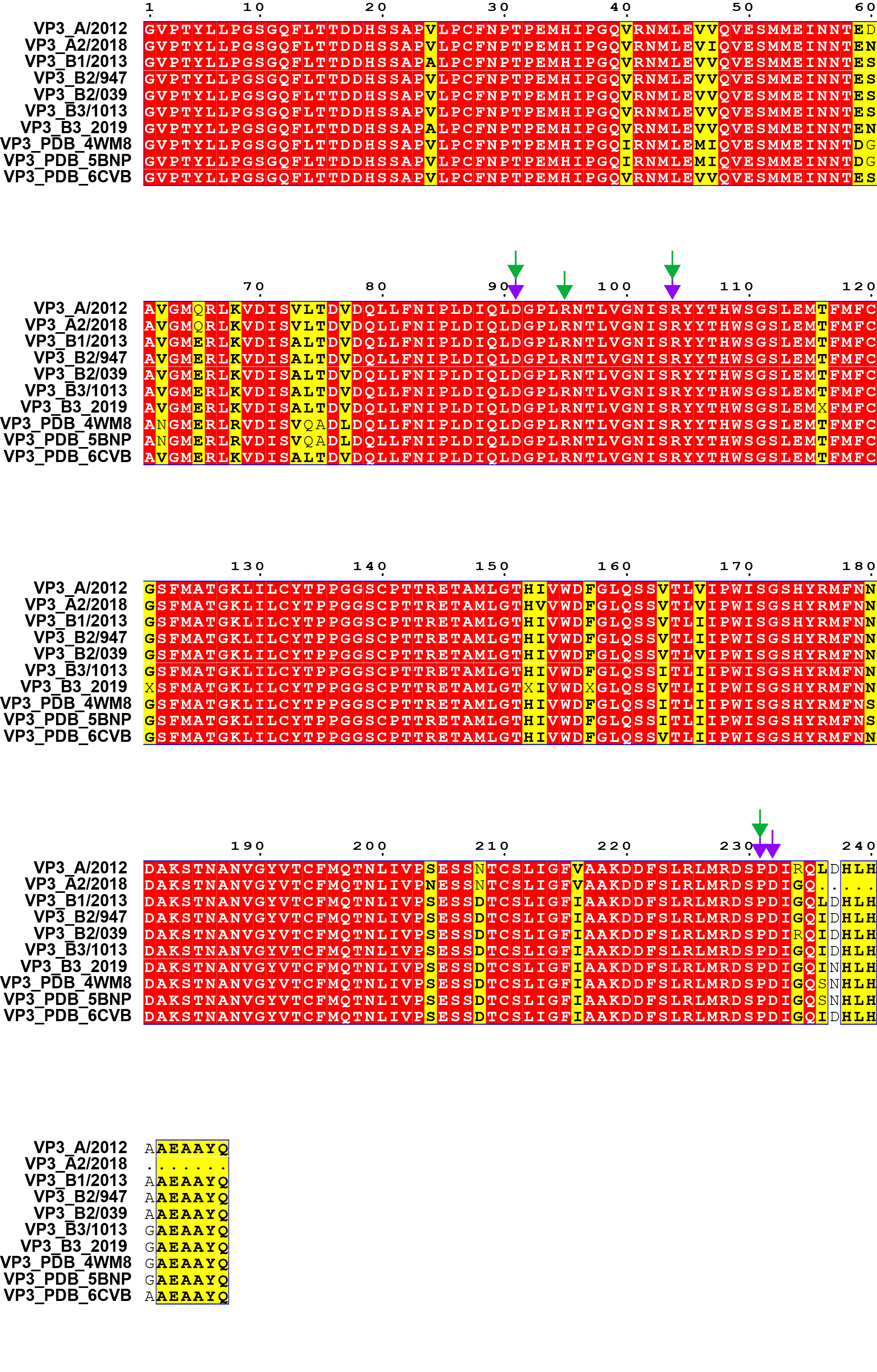

**Supplementary Figure 4. VP3 Multiple sequence alignment.** A multiple sequence alignment of the VP3 sequences of the sequenced Enterovirus-D68 virus stocks was performed with Clustal OMEGA^1^ . The ESPRIPT3.0 program^2^ was used to render sequence similarities. Purple (PDB: 5BNP) and green (PDB: 6CVB) arrows indicate residues in sialic acid coordination according to the Protein-Ligand Interaction Profiler PLIP^3^.

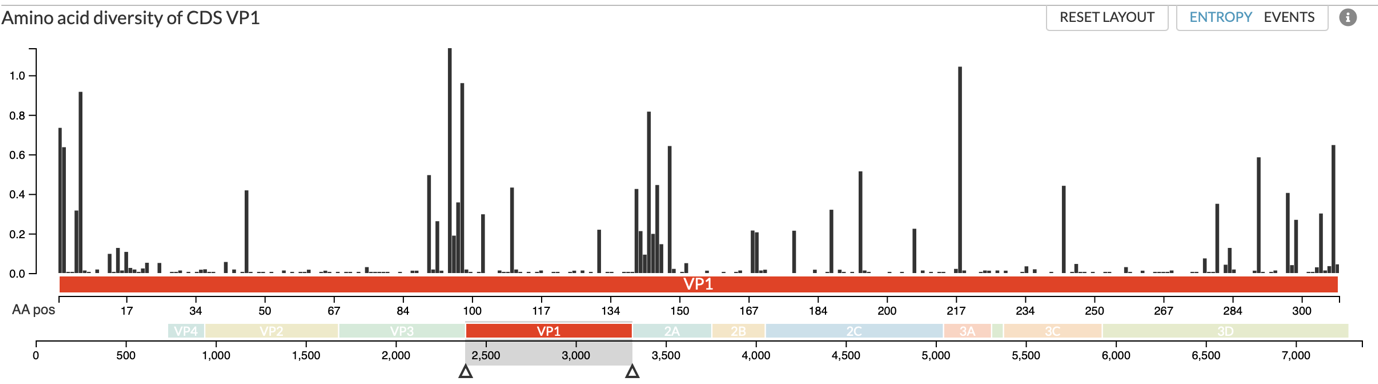
**Supplementary Figure 5. Nextstrain analysis of VP1 amino acid diversity.** The data displayed are derived from the Nextstrain^4^ platform focusing on amino acid polymorphisms in the non-structural protein VP1.

**Polymeric (2,8)-2,3-sialyl lactose**

**General materials and methods**

Recombinant enzymes PmST1 M144D^5^ , NmCSS^6^ and CSTII were expressed and purified according to reported protocols.All chemicals were obtained from commercial sources and were used without further purification unless described otherwise. Synthesis grade solvents were used and stored under argon over 4Å molecular sieves. TLC analysis was performed on SilicaPlate TLC Aluminum Backed TLC F254 (SiliCycle)plates. Spots were visualised with UV light (254 nm), 10% sulfuric acid in methanol and Ceric Ammonium Molybdate (4 ml H2SO4, 36 ml H2O, 1 g ammonium molybdate and 400 mg ceric ammonium sulfate). 1H NMR and 13C NMR spectra were recorded on an Agilent 400 spectrometer (300 MHz and 101 MHz) and Bruker Avance Neo 600 spectrometer (600 MHz). Chemical shifts are reported in ppm relative to TMS (0.00 ppm for 1H NMR). Spectra were recorded in D2O using the solvent as the internal standard in 1H NMR (D2O: 4.79 ppm 1H). Dialysis was done using dialysis tubing (MWCO:14kDa), benzoylated dialysis tubing (MWCO:2000) obtained from Sigma-Aldrich, SnakeSkinTM dialysis tubing (MWCO: 3.5kDa) or Slide-A-Lyser dialysis cassettes (MWCO: 10 kDa) obtained from Thermo-Fisher.

**Synthesis of S2**

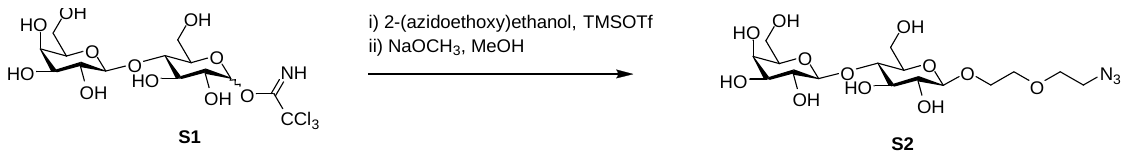

Lactoside **S1**^7^ (0.895 g, 1.2 mmol) and 2-(azidoethoxy)ethanol (0.614 g, 4.8 mmol, 4 eq.) were dissolved in dry DCM and stirred under an argon atmosphere. The reaction was cooled to 0°C and trimethylsilyl trifluoromethanesulfonate (0.065 mL, 0.35 mmol, 0.3 eq.) was added dropwise. The reaction was stirred until TLC indicated completion and quenched by dropwise addition of DI water. The mixture was washed thrice with DCM. Organic phases were combined and washed with DI and brine. Purification by silica gel chromatography (25% EtOAC in PE) to afford a white fluffy solid. The acetylated lactoside was dissolved in methanol and a catalytic amount of sodium methoxide was added. The mixture was stirred at room temperature for 2h. Upon completion, the reacton was neutralised with amberlite H+ resin. The resin was filtered off and the solvent was evaporated to provide product **S2** in 42% yield.

NMR was in accordance with previously described spectra^8^.

**Synthesis of S4**

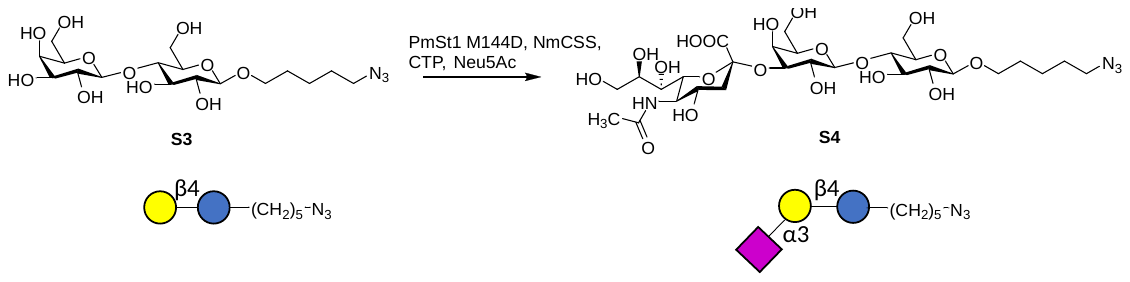

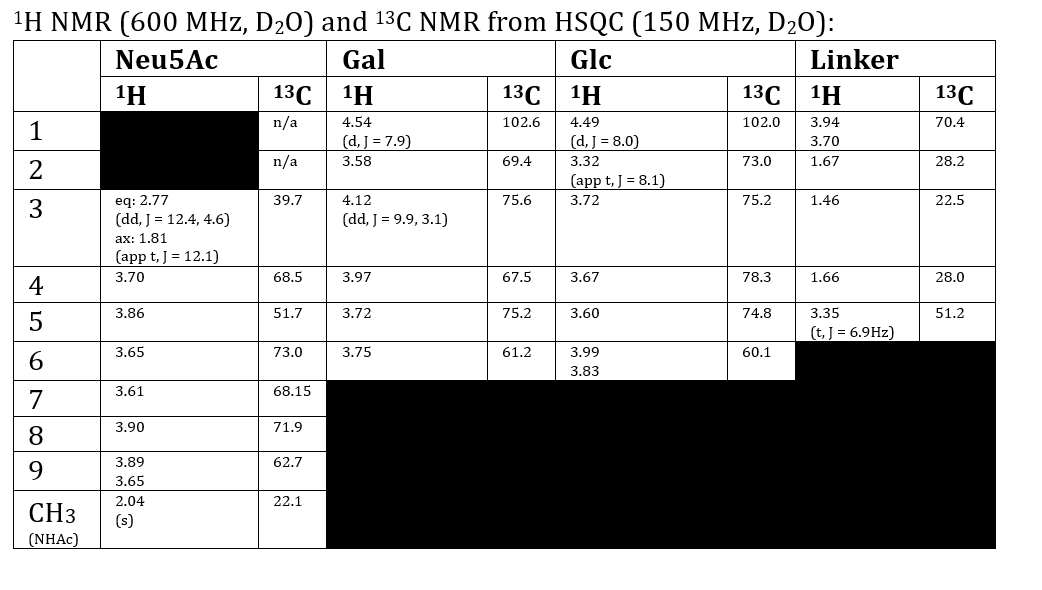
**S3**^9^ (50 mg,0.12 mmol ), Neu5Ac (75 mg, 0.24 mmol, 2 eq) and CTP (175 mg, 0,36 mmol, 3 eq) were dissolved in Tris-HCl buffer (100 mM, pH 7.2) to a concentration of 10 mM. Pmst1 M144D (0.01 mg/mL) and MnCSS (0.01 mg/ml) were added and the mixture was incubated at 37 °C until ESI-MS confirmed full conversion of starting material. Reaction mixture was lyophilized and purified through silica column chromatography to yield α2,3-sialyllactose **S4** as a white fluffy solid in 69% yield.

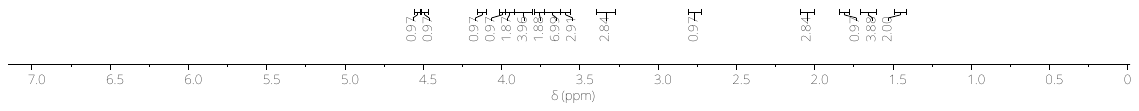

**Supplementary Figure 6.** 600 MHz ^1^H NMR spectra of S4

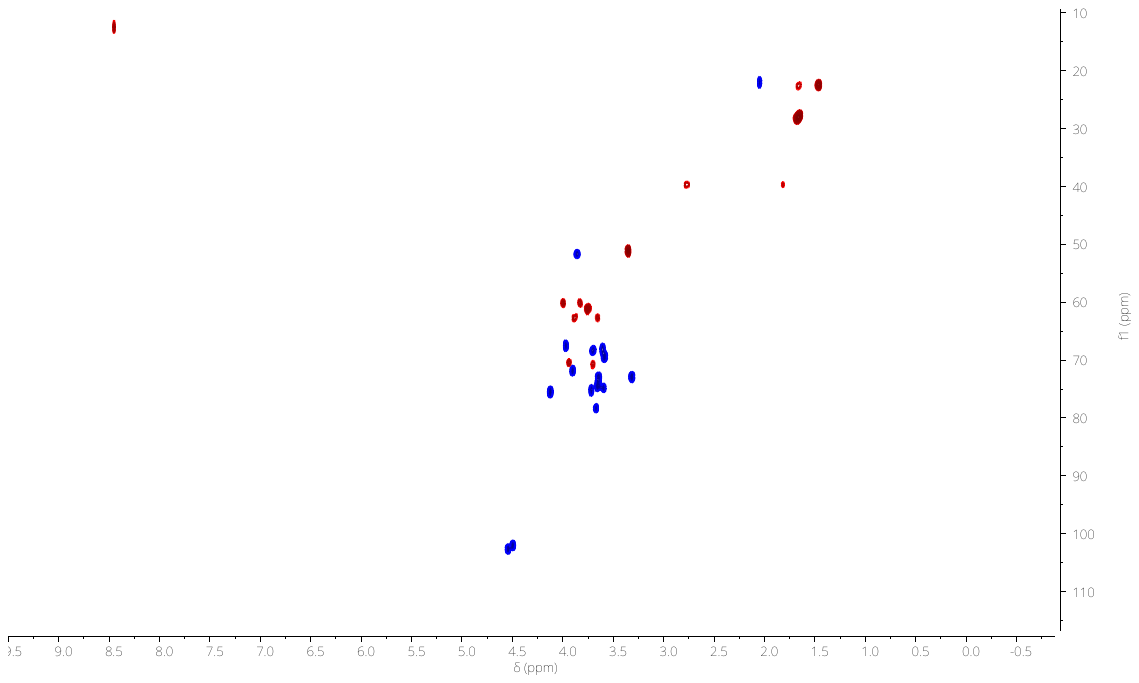

**Supplementary Figure 7.** 600 MHz ^13^C-^1^H HSQC NMR spectra of S4

**Synthesis of S5**

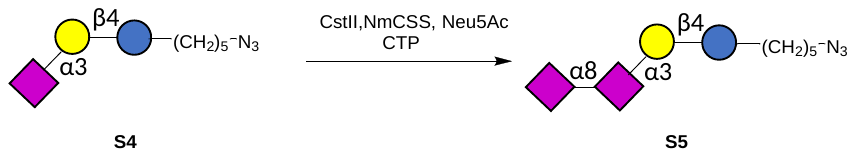

**S4** (42 mg,0.06 mmol), Neu5Ac (28 mg, 0.09 mmol, 1,5 eq) and CTP (63.5 mg, 0,12 mmol, 2 eq) were dissolved in Tris-HCl buffer (100 mM, pH 7.2) to a concentration of 10 mM. CstII (0.01 mg/mL) and MnCSS (0.01 mg/ml) were added and the mixture was incubated at 37 °C for 5h until ESI-MS confirmed maximal product formation. Reaction mixture was lyophilized and purified by size-exclusion chromatography (BioRad P-6, 45-90µm) with 0.05 M NH_4_HCO_3_ as eluent to yield α2,8-α2,3-sialyllactose **S5** in 51% yield.

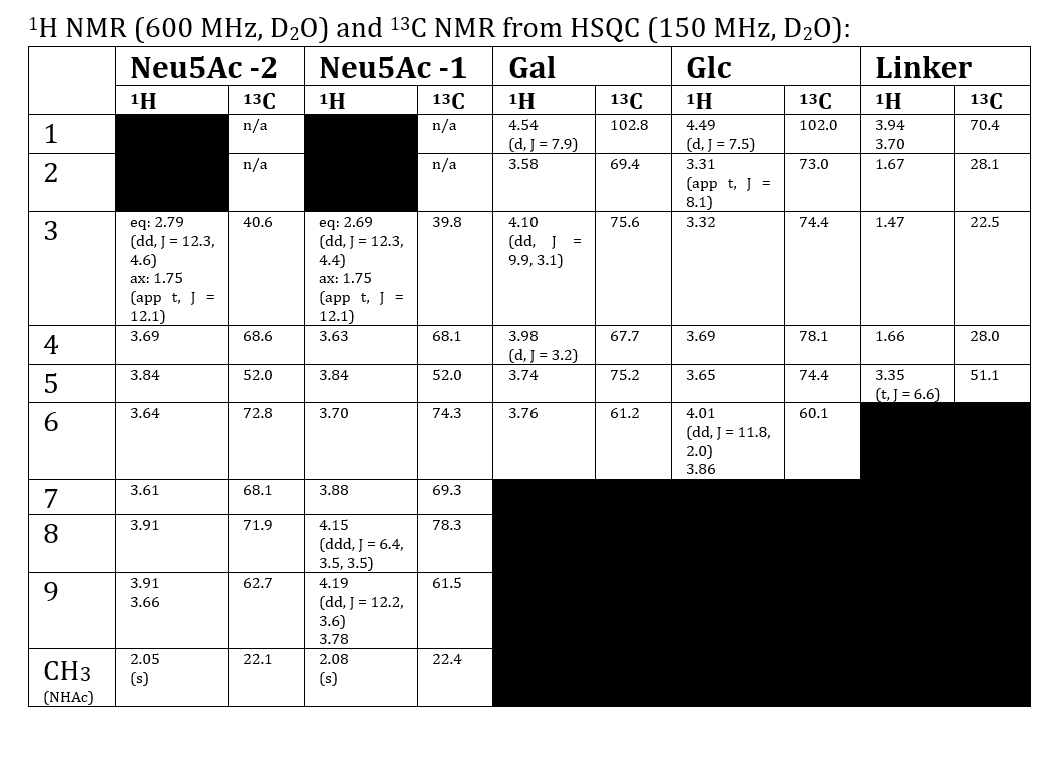

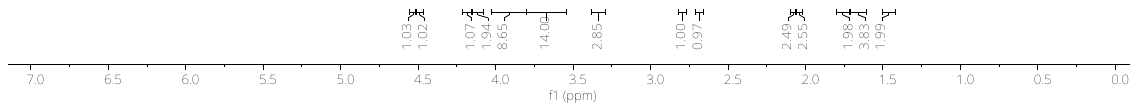

**Supplementary Figure 8.** 600 MHz ^1^H NMR spectra of S5

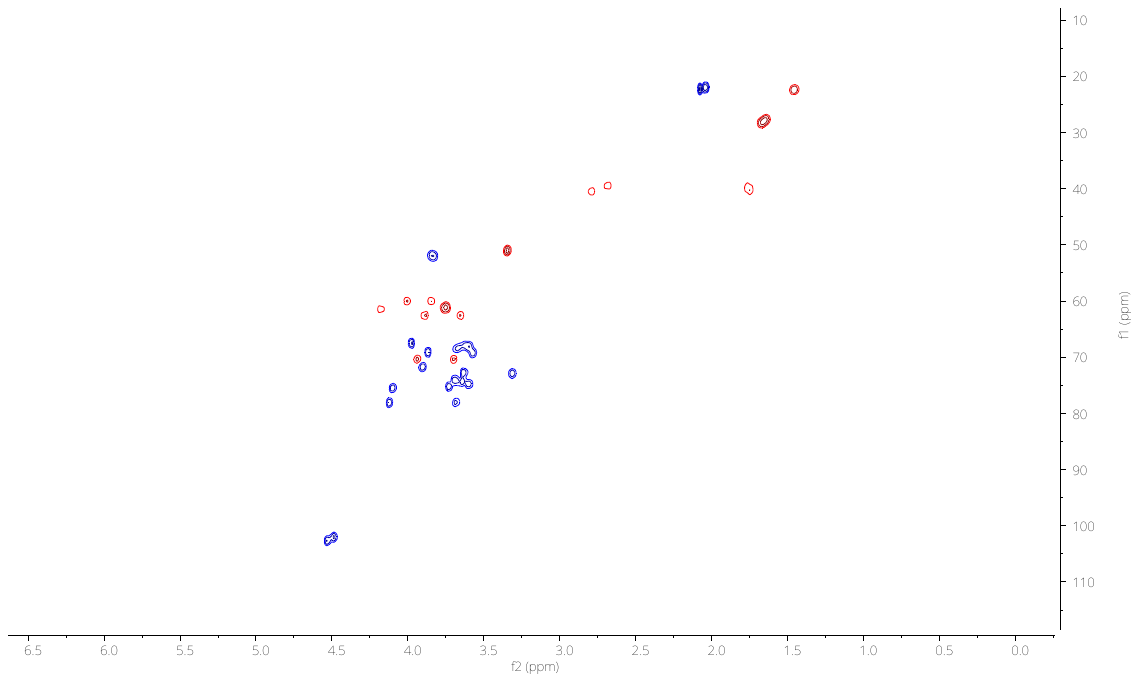

**Supplementary Figure 9:** 600 MHz ^13^C-^1^H HSQC NMR spectra of S5

**Hyperbranched Polyglycerol (hPG) synthesis**

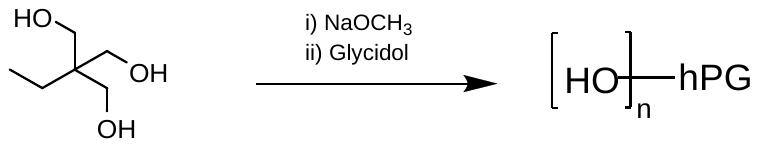

The polymerisation was carried out in a flask fitted with a mechanical stirrer and a syringe pump under argon atmosphere as described before^10^. 1,1,1- Tris(hydroxymethyl)propane (0.06 g, 1 equiv, 0.45 mmol) was added to the flask and melted at 60°C. Sodium methoxide solution (30% in MeOH, 16µl, 0.15 mmol,0.3 eq.) was added under nitrogen atmosphere and the resulting mixture was stirred for 30 min. Temperature of the reactor was increased to 95°C and 6 ml decane was added. Glycidol monomer (6ml) was added at 0.5 mL/hr using a syringe pump. After 16h, the reaction was quenched by addition of methanol. Reaction solvent was decanted and polymer was precipitated by the addition of acetone. Precipitate was redissolved in methanol and precipitation was repeated once more by addition of acetone(4eq/methanol). Precipitate was taken up in DI water and neutralised with drops 1M HCl and the crude compound was dialysed using a cellulose-based dialysis membrane (MWCO: 14kDa) against DI water for 72h, changing the water thrice daily. Product was lyophilized to give a colourless viscous polymer. Polymer size and degree of branching (DB) and monomer composition were calculated using ^1^H NMR and Inverse gated ^13^C NMR as described before^11^.

**Supplementary Table 1.** Characteristics of the hyperbranched polyglycerol

| Mn (kDa) | Degree of branching (DB) | 1,3-linear monomers (%) | 1,4-linear monomers (%) | Dendritic (%) | Terminal (%) |
| --- | --- | --- | --- | --- | --- |
| 14,9 | 0.57 | 10,4 | 25,4 | 27,9 | 36,3 |

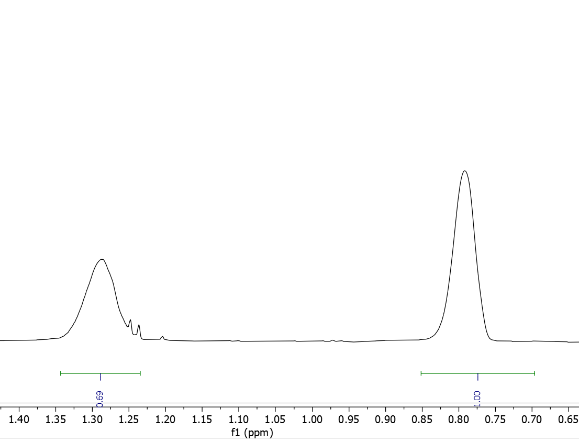

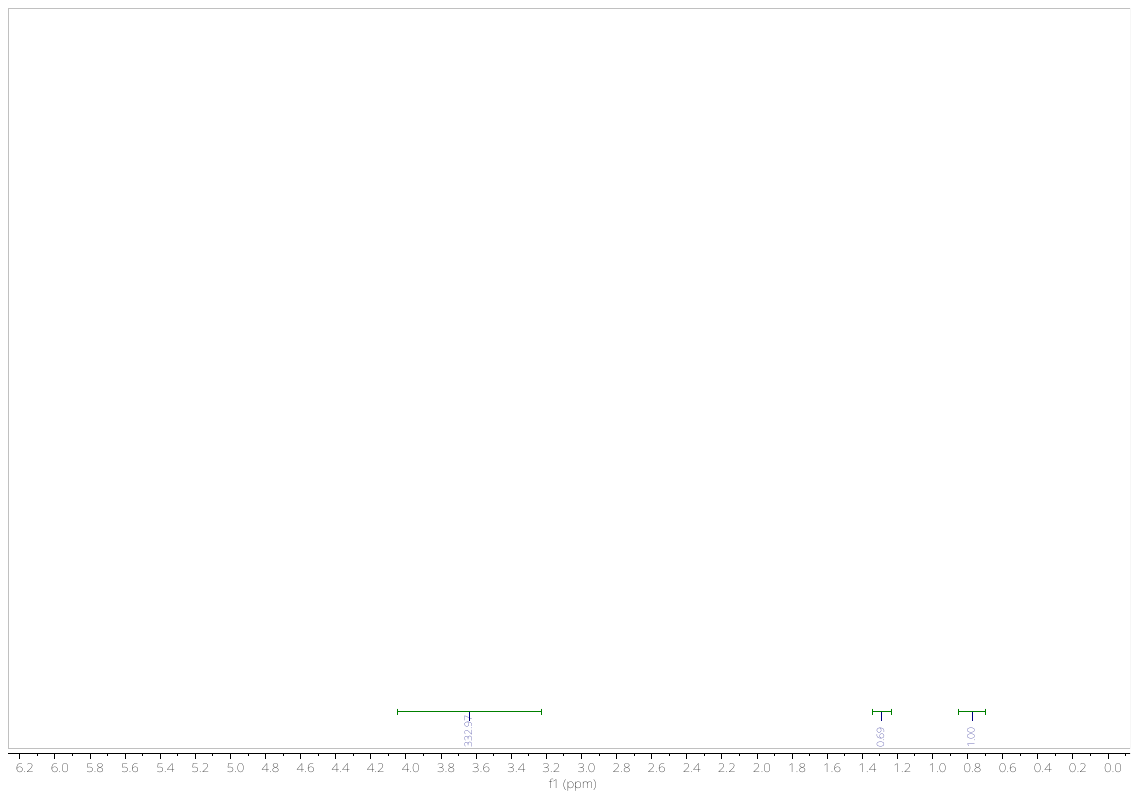

**Supplementary Figure 10.** 600 MHz ^1^H NMR spectra of hPG-OH

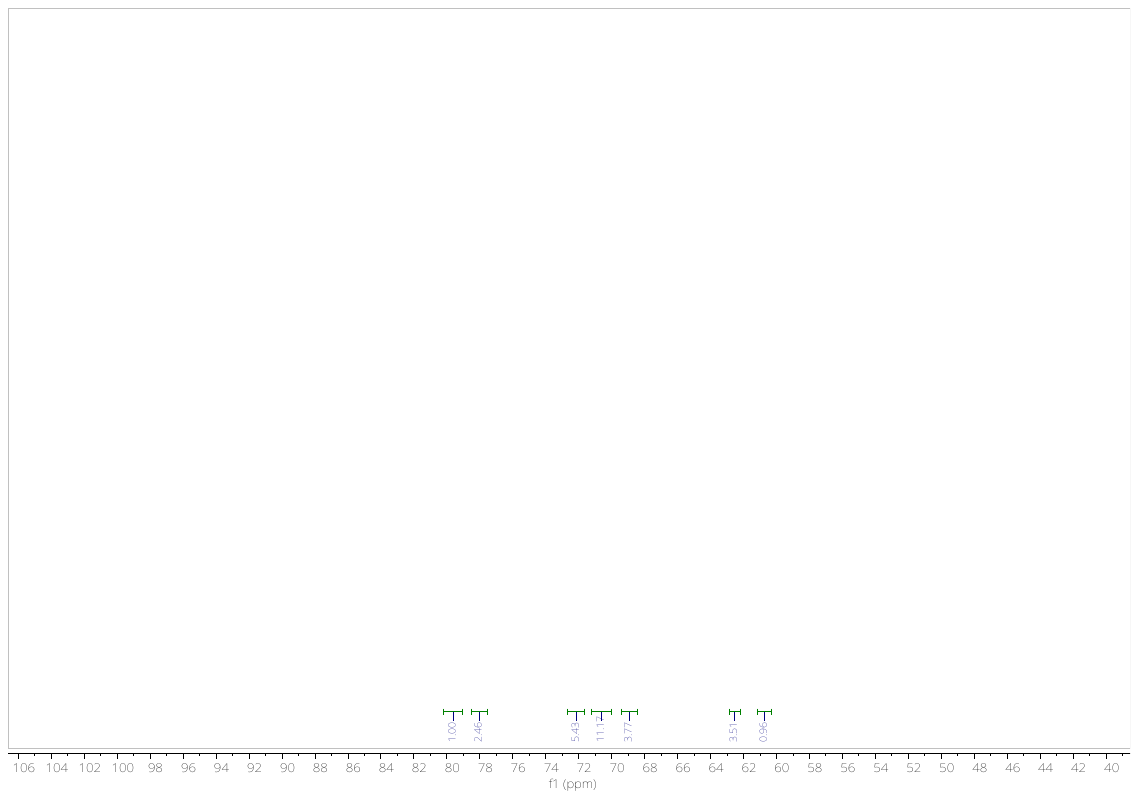

**Supplementary Figure 11.** 150 MHz ^13^C Inverse gated NMR spectra of hPG-OH

**Propargylation of hyperbranched polyglycerol**

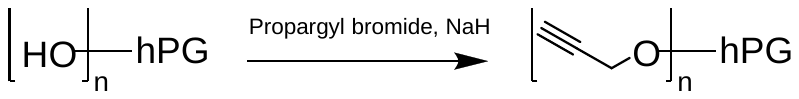

Number of open OH-groups in polymeric samples was determined by IG 13C NMR through determination of terminal and linear monomers in polymer as described before^11^. To a solution of hPG-OH in dry DMF, NaH (2.2 equiv, 60 % dispersion in mineral oil)was added under nitrogen atmosphere. After stirring for 3 h at r.t., potassium iodide ( 0.2 equiv) was added. Mixture was cooled to 0 °C , and propargyl bromide (1.5 equiv) was added dropwise and allowed to warm to r.t.. After stirring for 16h under nitrogen atmosphere, mixture was diluted with EtOAc. Organic layer was washed with DI water. After evaporation of EtOAc, crude product was redissolved in DCM and transferred to benzoylated dialysis tubing (MWCO: 2 kDa). Product underwent dialysis in DCM for 72 h, changing DCM thrice daily. Product was extracted from dialysis bag and solvent was evaporated under reduced pressure to obtain **hPG-alkyne** a brown viscous oil in 74% yield. Degree of functionalization was 62%, determined though previously described NMR integration method^11^.

^1^H NMR (400 MHz, CDCl_3_) δ 4.33 (ap. s., C*H*HCCH), 4.18 (ap. s., CH*H*CCH ), 4.06 – 3.24 (m, 500H, hPG backbone ), 2.49 (s, 62H).

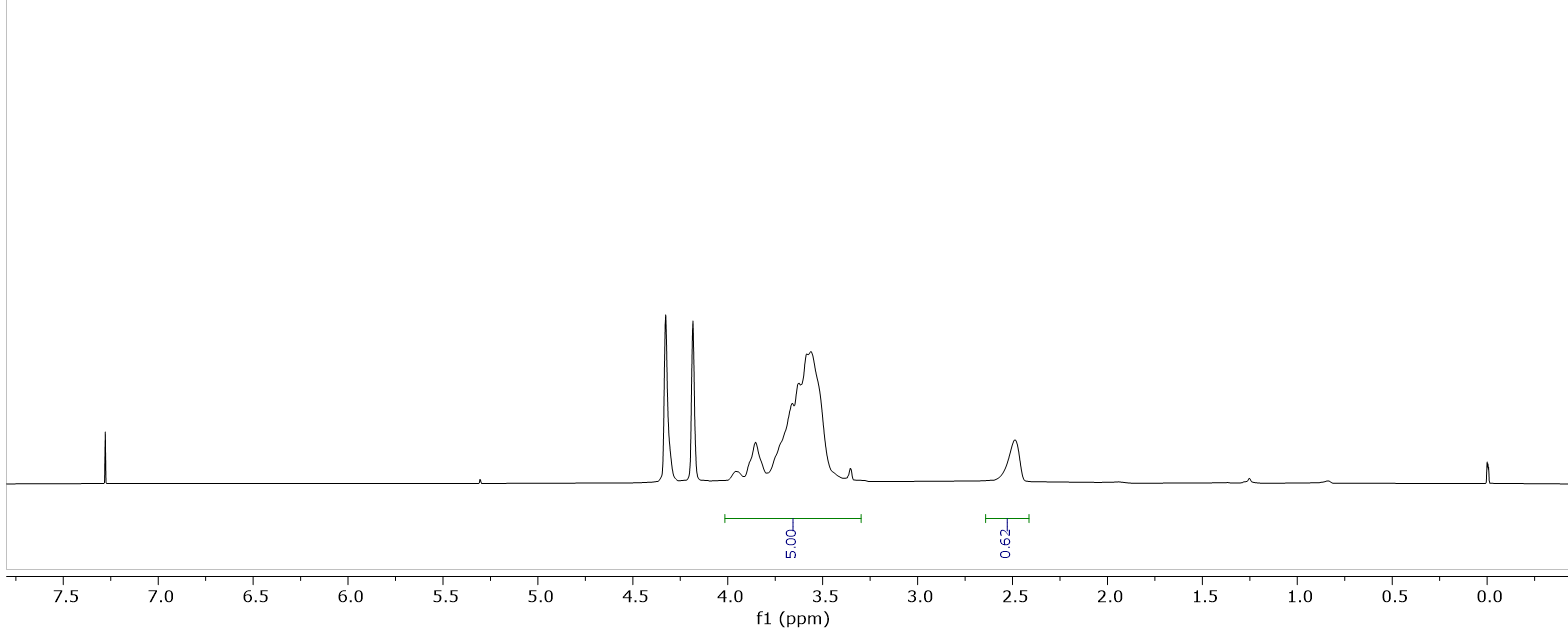

**Supplementary Figure 12.** 600 MHz ^1^H NMR spectra of hPG-propargyl

**hPG-2,3-sialyl lactose**

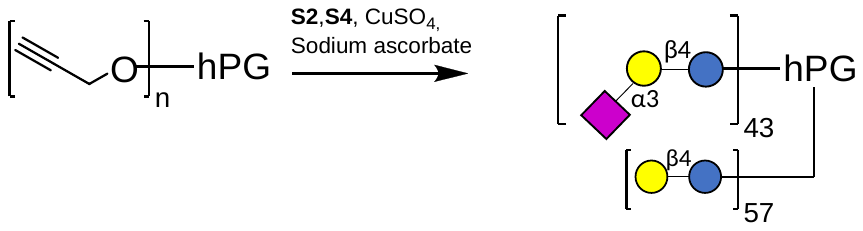

**hPG-alkyne** (1 mg, 6.7 µmol alkyne groups) was added to a vial from an 10 mg/ml aliquot solution. 2,3-α-sialyl-Lactose **S4** (2.8 mg, 4.0 µmol, 0.6 eq.) and **6** (1.8 mg, 4.0 µmol, 0.6 eq.) were dissolved in DMF separately and transferred to the vial. Sodium ascorbate (0.4 mg, 2.0 µmol, 0.3 eq.) and Copper (II) sulfate pentahydrate (0.17 mg, 0.67 µmol, 0.1 eq.) were dissolved in DI water separately and added to the vial from a freshy made aliquot solution of 10 mg/ml. Argon was bubbled through the solution for 5 min and the vial was sealed and allowed to stir for 7 days at room temperature. EDTA (0.5 mg, 1.34 µmol, 0.5 eq.) was added from a freshly made aliquot of 10 mg/ml and the mixture was allowed to stir for 2h. Reaction mixture was transferred to dialysis cassette (MWCO: 10kDa) and reaction vial was washed with DI water. Mixture was dialysed against DI water for 3 days and water was replaced thrice a day. Mixture was transferred to a centrifuge spinfilter tube (MWCO: 15kDa). Mixture was centrifuged and redissolved thrice with DI water, thrice with 0.05M ammonium bicarbonate and thrice with DI water. Residue is dissolved in DI water and lyophilised to obtain **hPG-2,3** as an white solid in 41% yield. Functionalization based on triazole and ligand signal was determined to be 43% **S4** and 57% **S2**.

^1^H NMR (600 MHz, D_2_O) δ 7.98 (bs, 1H, triazole), 4.53 (hPG), 4.45 (d, *J* = 7.8 Hz, H1 **S4**), 4.37 (d, *J* = 7.7 Hz, H1 **S4**, H1 and H1’ **S2**), 4.04 (ap. d, *J* = 10.0 Hz, H3 Gal **S4**), 3.94 – 3.40 (m, 33H, hPG and glycan protons), 3.25 – 3.22 (m, Linker C**H**_2_-O-**S4**/**S2**), 2.68 (dd, *J* = 12.6, 4.7 Hz, H3 eq **S4**), 1.95 (s, s, NHAc **S5**), 1.73 (m, H3 ax **S4** and Linker **S4**), 1.56 (m, Linker **S4**), 1.22 (m, Linker **S4**).

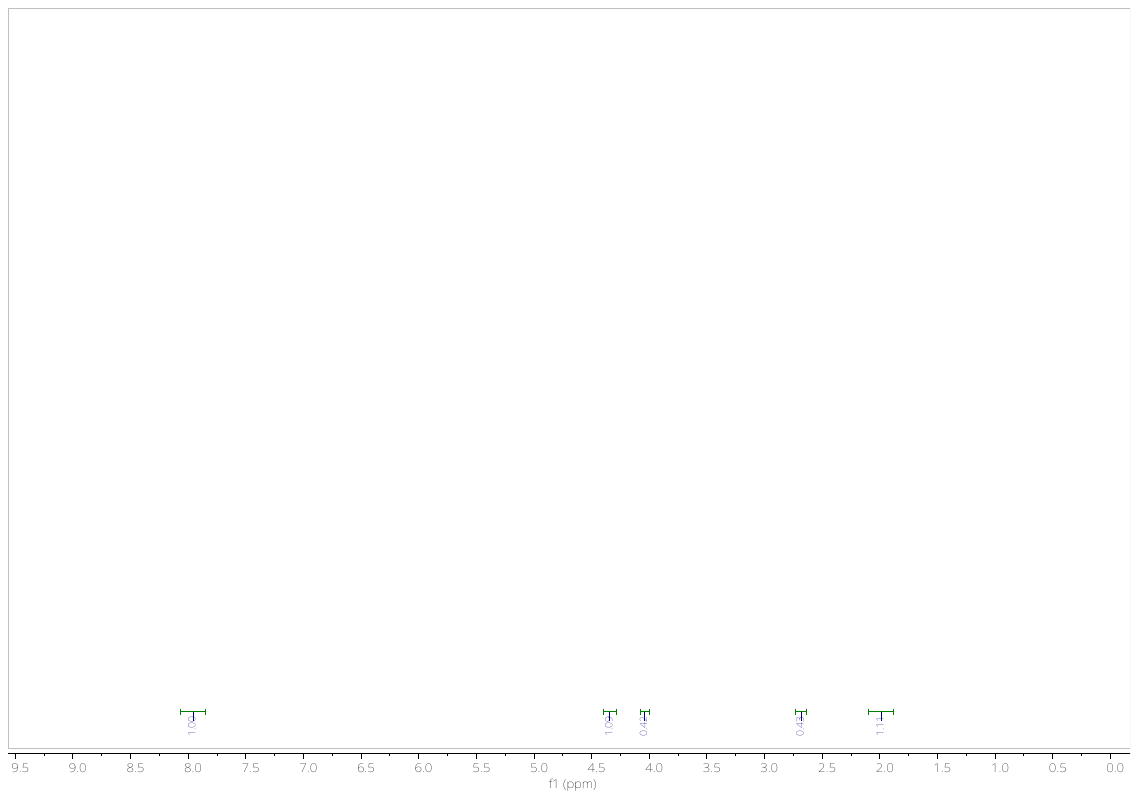

**Supplementary Figure 12.** 600 MHz ^1^H NMR spectra of hPG-2,3

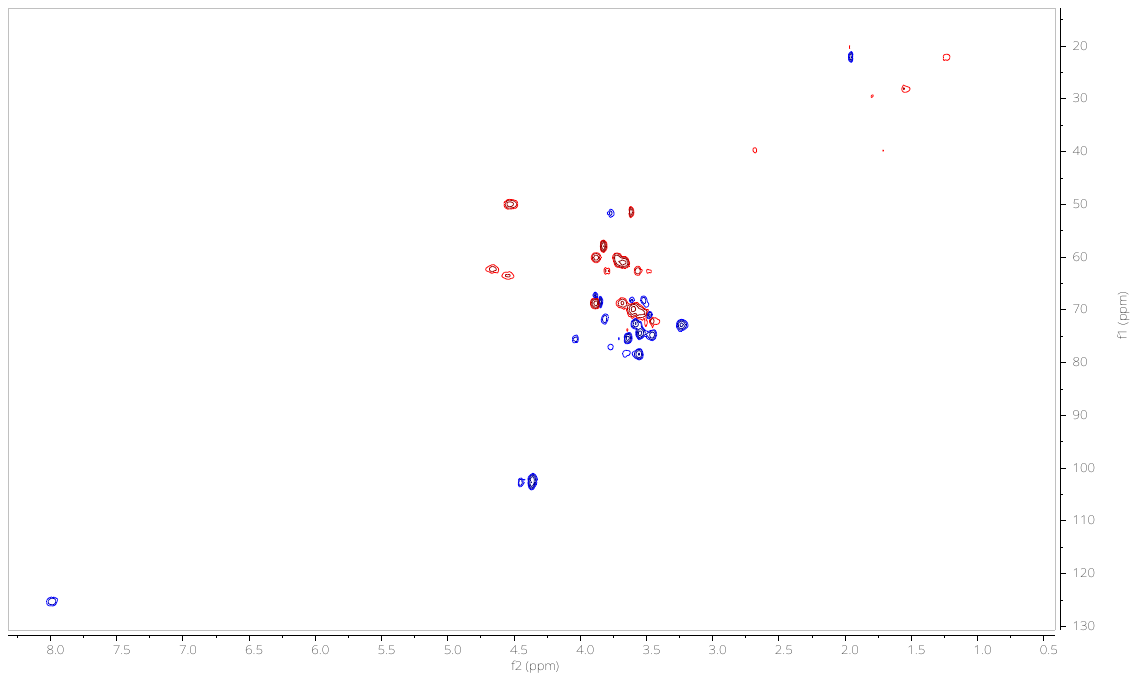

**Supplementary Figure 13.** 600 MHz ^13^C-^1^H HSQC NMR spectra of hPG-2,3

**hPG-2,3-2,8-sialyl lactose**

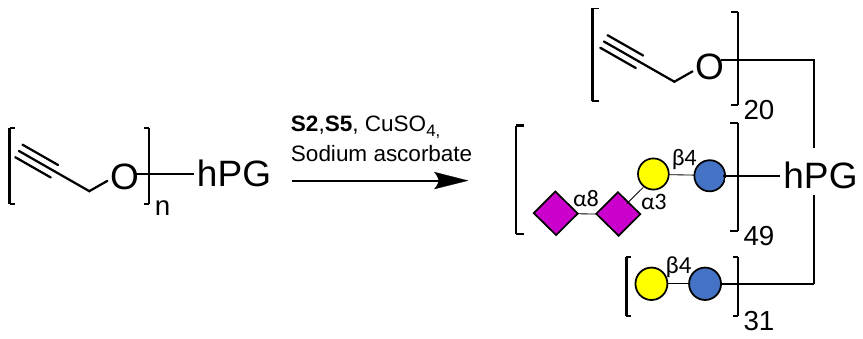

hPG-alkyne (1 mg, 6.7 µmol alkyne groups) was added to a vial from an 10 mg/ml aliquot solution. 2,8-α-sialyl-2,3-α-sialyl-Lactose **S5** (4.2 mg, 4.0 µmol, 0.6 eq.) and **S2** (1.8 mg, 4.0 µmol, 0.6 eq.) were dissolved in DMF separately and transferred to the vial. Sodium ascorbate (0.4 mg, 2.0 µmol, 0.3 eq.) and Copper (II) sulfate pentahydrate (0.17 mg, 0.67 µmol, 0.1 eq.) were dissolved in DI water separately and added to the vial from a freshy made aliquot solution of 10 mg/ml. Argon was bubbled through the solution for 5 min and the vial was sealed and allowed to stir for 7 days at room temperature. EDTA (0.5 mg, 1.34 µmol, 0.5 eq.) was added from a freshly made aliquot of 10 mg/ml and the mixture was allowed to stir for 2h. Reaction mixture was transferred to dialysis cassette (MWCO: 10kDa) and reaction vial was washed with DI water. Mixture was dialysed against DI water for 3 days and water was replaced thrice a day. Mixture was transferred to a centrifuge spinfilter tube (MWCO: 15kDa). Mixture was centrifuged and redissolved thrice with DI water, thrice with 0.1M ammonium bicarbonate and thrice with Di water to remove unreacted ligand. Residue is dissolved in DI water and lyophilised to obtain **hPG-2,3-2,8** as an white solid in 32% yield. Functionalization based on triazole and ligand signal was determined to be 49% **S5,** 31% **S2** and 20% alkyne remaining.

^1^H NMR (600 MHz, D_2_O) δ 7.99 (s, 1H, triazole), 4.54 (hPG), 4.45 (d, *J* = 8.2 Hz, **S5**), 4.40 (d, *J* = 7.8 Hz, **S5**), 4.37 (d, *J* = 7.5 Hz, **S2**), 4.30 (ap. s, alkyne hPG), 4.19 (ap. s, alkyne hPG), 4.09 (dd, *J* = 12.4, 3.2 Hz, H3 Gal **S5**), 4.02 (m, H8,H9 Neu5Ac1 **S5**), 3.95 – 3.44 (m, 48H, hpG and ligand protons), 3.29 – 3.20 (m, 2H), 2.89 (s, alkyne-HPG), 2.71 (dd, *J* = 12.6, 4.6 Hz, H3 eq. Neu5Ac2 **S5**), 2.60 (dd, *J* = 12.5, 4.3 Hz, H3 eq. Neu5Ac1 **S5**), 2.01 – 1.98 (s, NHAc Neu5Ac1 **S5**), 1.97 – 1.93 (s, NHAc Neu5Ac2 **S5**), 1.68 (m, H3 ax **S4** and Linker **S4**), 1.57 (m, Linker **S4**), 1.37 (m, Linker **S4**).

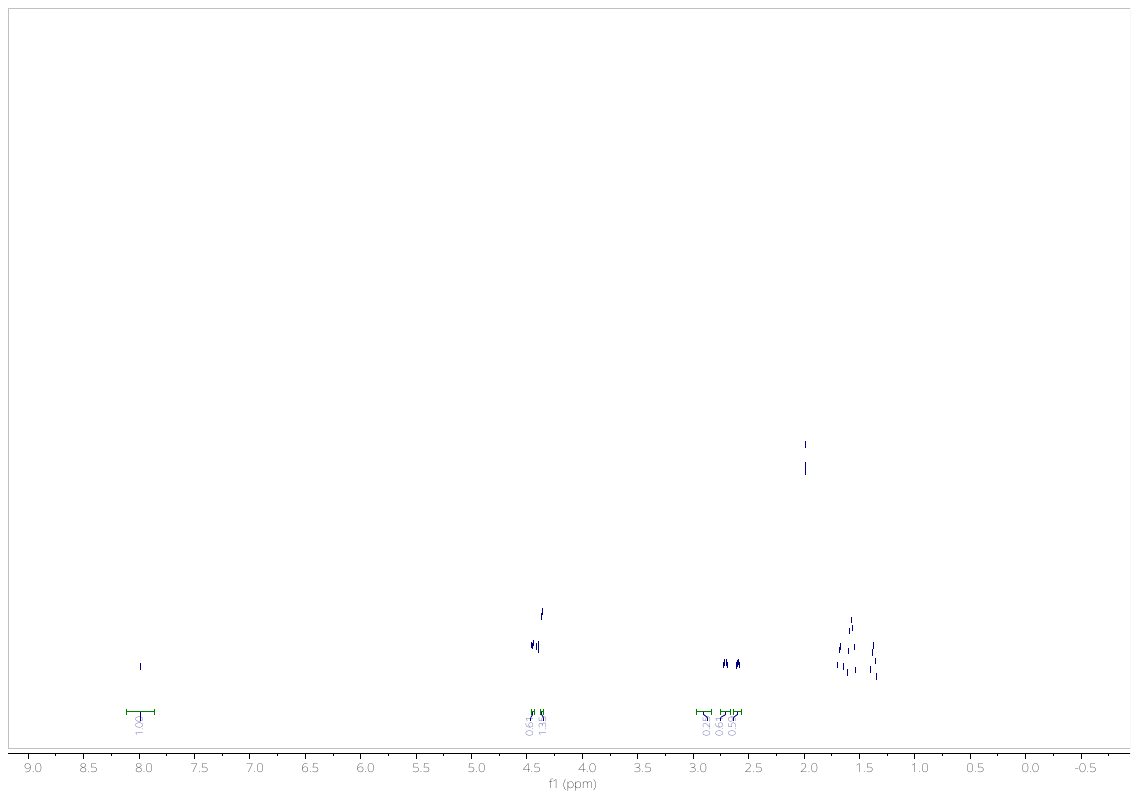

**Supplementary Figure 14.** 600 MHz ^1^H NMR spectra of hPG-2,3-2,8

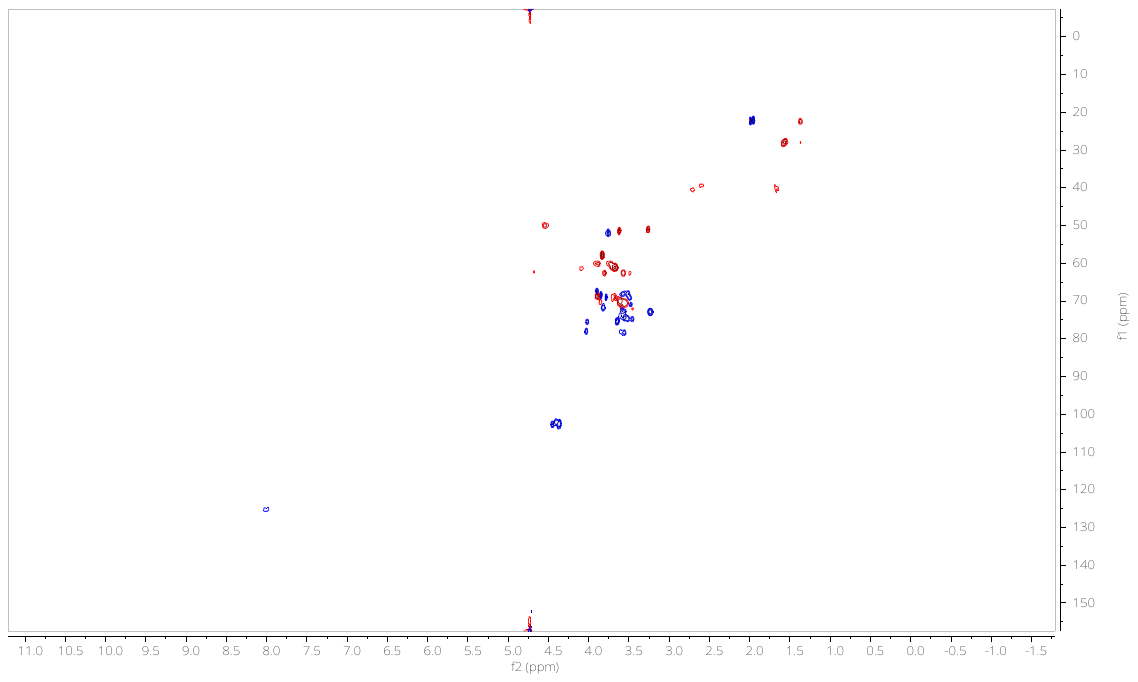

**Supplementary Figure 15.** 600 MHz ^13^C-^1^H HSQC NMR spectra of hPG-2,3-2,8

**References:**

(1)	Madeira, F.; Madhusoodanan, N.; Lee, J.; Eusebi, A.; Niewielska, A.; Tivey, A. R. N.; Lopez, R.; Butcher, S. The EMBL-EBI Job Dispatcher Sequence Analysis Tools Framework in 2024. *Nucleic Acids Res* **2024**, *52* (W1), W521–W525. https://doi.org/10.1093/nar/gkae241.

(2)	Robert, X.; Gouet, P. Deciphering Key Features in Protein Structures with the New ENDscript Server. *Nucleic Acids Research* **2014**, *42* (W1), W320–W324. https://doi.org/10.1093/nar/gku316.

(3)	*PLIP: fully automated protein–ligand interaction profiler | Nucleic Acids Research | Oxford Academic*. https://academic.oup.com/nar/article/43/W1/W443/2467865?login=false (accessed 2024-10-15).

(4)	*Nextstrain: real-time tracking of pathogen evolution - PubMed*. https://pubmed.ncbi.nlm.nih.gov/29790939/ (accessed 2024-10-21).

(5)	Sugiarto, G.; Lau, K.; Qu, J.; Li, Y.; Lim, S.; Mu, S.; Ames, J. B.; Fisher, A. J.; Chen, X. A Sialyltransferase Mutant with Decreased Donor Hydrolysis and Reduced Sialidase Activities for Directly Sialylating LewisX. *ACS Chem Biol* **2012**, *7* (7), 1232–1240. https://doi.org/10.1021/cb300125k.

(6)	Yu, H.; Yu, H.; Karpel, R.; Chen, X. Chemoenzymatic Synthesis of CMP-Sialic Acid Derivatives by a One-Pot Two-Enzyme System: Comparison of Substrate Flexibility of Three Microbial CMP-Sialic Acid Synthetases. *Bioorg Med Chem* **2004**, *12* (24), 6427–6435. https://doi.org/10.1016/j.bmc.2004.09.030.

(7)	Xue, M.; Tan, L.; Zhang, S.; Wang, J.-N.; Mi, X.; Si, W.; Qiao, Y.; Lao, Z.; Meng, X.; Yang, Y. Chemoenzymatic Synthesis of Sialyl-Α2,3-Lactoside–Functionalized BSA Conjugate Inhibits Influenza Infection. *European Journal of Medicinal Chemistry* **2024**, *276*, 116633. https://doi.org/10.1016/j.ejmech.2024.116633.

(8)	Naicker, K. P.; Li, H.; Heredia, A.; Song, H.; Wang, L.-X. Design and Synthesis of αGal-Conjugated Peptide T20 as Novel Antiviral Agent for HIV-Immunotargeting. *Org. Biomol. Chem.* **2004**, *2* (5), 660–664. https://doi.org/10.1039/B313844E.

(9)	Ledin, P. A.; Kolishetti, N.; Hudlikar, M. S.; Boons, G.-J. Exploring Strain-Promoted 1,3-Dipolar Cycloadditions of End Functionalized Polymers. *Chemistry – A European Journal* **2014**, *20* (28), 8753–8760. https://doi.org/10.1002/chem.201402225.

(10)	Ul-Haq, M. I.; Shenoi, R. A.; Brooks, D. E.; Kizhakkedathu, J. N. Solvent-Assisted Anionic Ring Opening Polymerization of Glycidol: Toward Medium and High Molecular Weight Hyperbranched Polyglycerols. *Journal of Polymer Science Part A: Polymer Chemistry* **2013**, *51* (12), 2614–2621. https://doi.org/10.1002/pola.26649.

(11)	Haksar, D.; Asadpoor, M.; Heise, T.; Shi, J.; Braber, S.; Folkerts, G.; Ballell, L.; Rodrigues, J.; Pieters, R. J. Fighting Shigella by Blocking Its Disease-Causing Toxin. *J. Med. Chem.* **2021**, *64* (9), 6059–6069. https://doi.org/10.1021/acs.jmedchem.1c00152.
